# Lung Tumour Cell-derived Exosomes Promote the Expansion of MDSC Through miR-21a/PDCD4/IL-6/STAT3 Pathway

**DOI:** 10.1101/708826

**Authors:** Xingju Zhang, Fei Li, Jiale Zhang, Ying Tang, Bin Xiao, Ying Wan, Shan Jiang

## Abstract

The percentage of MDSC population in peripheral blood is significantly increased in lung cancer patients. The accumulation of MDSC contributes to the development of a tumour-associated immune suppressive environment. But the mechanism mediating MDSC accumulation in lung cancer patients is elusive. We found that exosomes from Lewis lung cancer cells (LLC-Exo) can be taken up by the mice bone marrow cells and then promote MDSC expansion, which enhance tumour growth. Mechanistically, during the induction of MDSC from the bone marrow cells, miR-21a from LLC-Exo can stimulate BM cells to produce more IL-6 by inhibiting *pdcd4* expression post-transcriptionally. Autocrine IL-6 induces tyrosine 705(Y705) of STAT3 to be phosphorylated during MDSC expansion. LLC-Exo with miR-21a deletion lost its ability to stimulate IL-6 production, STAT3 activation and MDSC expansion. These results demonstrate that lung cancer cell-derived exosomes promote functional MDSC expansion through activation of miR-21a/PDCD4/IL-6/STAT3 pathway.

## Introduction

Myeloid-derived suppressor cells (MDSC) are a heterogeneous population that is comprised of immature myeloid cells generating in the bone marrow with the ability to suppress T cell activation(*D. I. Gabrilovich and Nagaraj, 2009; Makarenkova et al., 2006; Ostrand-Rosenberg and Sinha, 2009*). In mice, MDSC are characterised by expressing Ly-6C/G and CD11b molecules simultaneously(*D. I. Gabrilovich and Nagaraj, 2009; Ostrand-Rosenberg and Sinha, 2009*), while in humans MDSC are defined by the CD11b^+^CD33^+^HLA-DR^−^ phenotype(*Filipazzi et al., 2007; Mirza et al., 2006*). Abnormal expansion of MDSC is an important mechanism for tumour immune evasion and plays a causal role in tumour development(*D. I. Gabrilovich and Nagaraj, 2009; Ostrand-Rosenberg and Sinha, 2009*). In patients with bladder carcinoma, multiple myeloma, breast cancer and renal cell carcinoma, accumulation of MDSC was detected in peripheral blood (*Brimnes et al., 2010; Diaz-Montero et al., 2009; Ochoa et al., 2007; Yuan et al., 2011*). Tumour cells produce various factors including cyclooxygenase-2 (COX2)(*Rodriguez et al., 2005*), prostaglandins(*Sinha et al., 2007*), stem-cell factor (SCF)(*Pan et al., 2008*), macrophage colony-stimulating factor (M-CSF)(*Menetrier-Caux et al., 1998*), IL-6(*Menetrier-Caux et al., 1998; C. T. Wu et al., 2012; L. Wu et al., 2017*), granulocyte/macrophage colony-stimulating factor (GM-CSF)(*Serafini et al., 2004*) and vascular endothelial growth factor (VEGF)(*D. Gabrilovich et al., 1998*) to promote the expansion of MDSC through stimulation of myelopoiesis and inhibiting the differentiation of mature myeloid cells. Studies indicate that STAT3 is arguably the main transcription factor regulating the expansion of MDSC(*Guo et al., 2018a; Kortylewski et al., 2005; L. Li et al., 2014; Liu et al., 2010; Nefedova et al., 2005*).

Exosomes are endocytic originated small membrane vesicles (30 to 100 nm in diameter) and secreted by both normal and malignant cells(*Bobrie and Thery, 2013; Thery, 2011*). Exosomes can shuttle proteins and fragments of DNA, RNA and microRNA from donor to recipient cells(*Thery, 2011; Valadi et al., 2007*). Tumour-derived exosomes (TEX) play an important role in enhancing tumour development by inhibiting immune cell function (T and NK cells)(*Ashiru et al., 2010; Muller et al., 2016*), dentritic cell differentiation(*Zhou et al., 2014*) and stimulating regulatory cell formation (Tregs and Bregs) (*Y. Li et al., 2015; Wieckowski et al., 2009*). TEX can also induce MDSC expansion(*Chen et al., 2017; Guo et al., 2018a; Guo et al., 2018b; Liu et al., 2015; Liu et al., 2010; Xiang et al., 2009*). Furthermore, TEX promoted MDSC accumulation through tumour exosomal prostaglandin E2 (PGE2) and TGF-β molecules in murine mammary adenocarcinomas cells(*Xiang et al., 2009*). Heat shock protein 72 (HSP72) expressed at the surface of TEXs can have the effect of triggering MDSC immunosuppressive function via STAT3 activation in murine thymoma cells, mammary carcinoma cells and colon carcinoma cells(*Chalmin et al., 2010*). miR-10a, miR-21a, miR-29a and miR-92a in TEX from glioblastoma cells are responsible for the expansion and immunosuppressive function of MDSC(*Guo et al., 2018a; Guo et al., 2018b*). The percentage of MDSC population in peripheral blood is reported to be significantly increased in lung cancer patients when compared to that in healthy people and is negatively correlated with the time of progression-free survival in lung cancer patients(*de Goeje et al., 2015; Heuvers et al., 2013; Vetsika et al., 2014*). Circulating MDSC in NSCLC patients were predicted to be a marker for poor response to chemotherapy(*Huang et al., 2013; Vetsika et al., 2014*) and these cells were found to have strong suppressive functions to T cells activation(*Heuvers et al., 2013; Huang et al., 2013*). This means that MDSC in lung cancer patient exerts negative regulation on immune cells (DC/monocyte/T cells) and contributes to the development of a tumour-associated immune suppressive environment(*Vetsika et al., 2014*). There is little known, however, about the mechanism mediating significant accumulation of MDSC in lung cancer patients.

In this study, we reported that exosomes from mice Lewis lung cancer cells (LLC-Exo) taken up by mice bone marrow cells can promote MDSC expansion and enhance tumour growth significantly. Mechanistically we found that miR-21a molecule which is abundant in TEXs could be delivered to mice BM cells through phagocytosis of LLC-Exo and miR-21a stimulated BM cells to produce more IL-6 by inhibiting *pdcd4* expression post-transcriptionally. The autocrine IL-6 then induces more tyrosine 705(Y705) of STAT3 to be phosphorylated in BM cells and finally results in MDSC expansion. The LLC-Exo with miR-21a deletion lost its ability to stimulate IL-6 production and STAT3 activation in BM cells and could not promote MDSC expansion. Overall, we considered LLC-Exo to be able to promote functional MDSC expansion via the miR-21a/PDCD4/IL-6/STAT3 signalling pathway.

## Results

### TEX from LLC cells promote tumour growth and result in MDSC systemic accumulation in a tumour-bearing mouse

To determine whether TEX from lung cancer cells are responsible for tumour development in the patient, we isolated TEX from the supernatant of the culture medium with Lewis lung cancer cells (LLC-Exo) and exosomes from the medium cultured with normal fibroblast 3T3 cells (3T3-Exo) (Supplementary Fig. 1A, B). LLC-Exo and 3T3-Exo mixed with Lewis lung cancer cells (LLC) expressing luciferase were respectively injected into the mouse subcutaneously. The tumour growth in each mouse was monitored by the luminescent flux of tumour cells and total tumour volume (Supplementary Fig. 1C). On the fifteenth day the mice were sacrificed and the percentage of MDSCs (CD11b^+^Gr-1^+^) in the spleen of each mouse was calculated by FACS. In mice with an LLC-Exo injection the tumours formed faster and had significantly accelerated growth compared with the tumours in the control mice with 3T3-Exo injection (Fig. 1A, B; Supplementary Fig. 1D; n=8). We saw that MDSC population (CD11b^+^ Gr-1^+^) significantly increased in the spleens of mice with LLC-Exo injection compared to those from the control mice with 3T3-Exo injection (Fig.1C).

**Figure 1.**
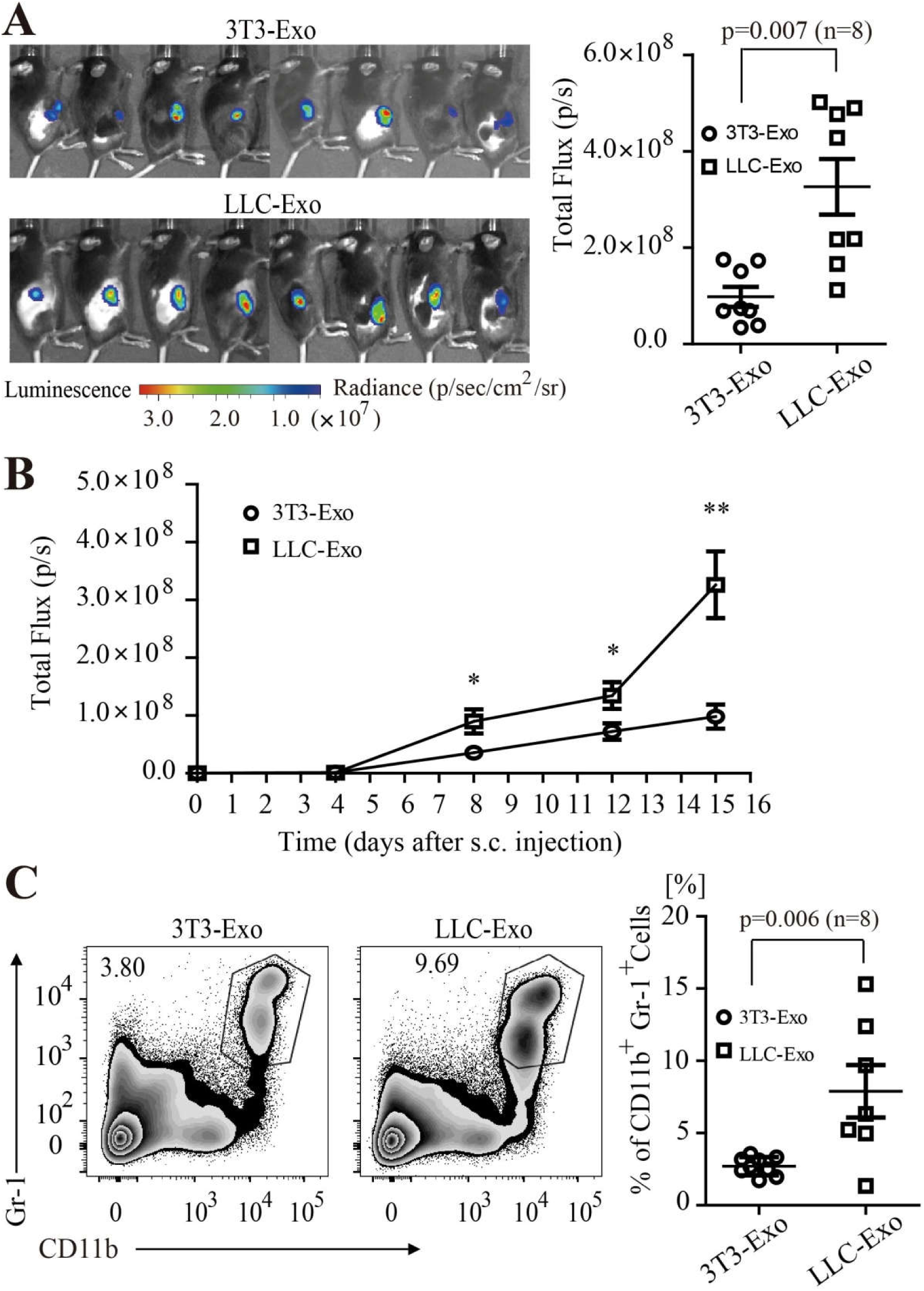
LLC-Exo promote tumour growth in mice. LLC cells with luciferase expression (1×10^6^ cells in 100 μL PBS/ mouse) mixed with exosomes (100 μg in 100 μL PBS /mouse) produced by 3T3 cells (3T3-Exo) or LLC cells (LLC-Exo) were subcutaneously injected into the C57BL/6 mouse. Each group consisted of eight animals. (**A**) An image of luminescence in the tumour of the mice at day15^th^ of injection. (**B**) Tumour growth in the mice was monitored every three days by using IVIS imager and the total luminescent fluxes of tumours in each mouse were calculated. * p<0.05; ** p<0.01, student t test. (**C**) One day after the last imaging process, the spleens were harvested. The percentage of MDSC (CD11b^+^Gr-1^+^) was analysed by FACS. The data is representative of two independent experiments. Error bars represent the mean ± sem.

The above results indicate that TEX from LLC cells promotes tumour growth dramatically and results in accumulation of MDSC in tumour bearing mice. Studies report that TEX from several different types of tumour cells could induce MDSC expansion (*Chen et al., 2017; Guo et al., 2018a; Guo et al., 2018b; Liu et al., 2015; Liu et al., 2010; Xiang et al., 2009*). All this made us curious about the possibility that LLC cells benefit their growth in mice via LLC-Exo to induce MDSC expansion. The MDSC are a heterogeneous population that is comprised of progenitors and precursors of myeloid cells. A block of immature myeloid cells (IMC) differentiation into mature myeloid cells in bone marrow results in an expansion of MDSC(*D. I. Gabrilovich and Nagaraj*). Studies also indicated that TEX could be taken up by the bone marrow cells(*Yu et al., 2007*). Consequently, we labeled LLC-Exo with PKH67 fluorescent dye and injected the mouse through a tail vein. Twenty-four hours later, there was an obvious portion of BM cells and splenocytes that proved PKH67-positive after flow cytometry analysis (Supplementary Fig. 2A, B). We also cultured PKH67-labled LLC-Exo with mice BM cells for four to five hours in-vitro and found LLC-Exos can be phagocytosed by mice BM cells through confocal microscopy analysis (Supplementary Fig. 2C).We considered these LLC-Exo which were taken up by BM cells probably stimulate MDSC to expansion through a certain mechanism.

### LLC-Exo promote MDSCs expansion by stimulating IL-6 production in BM cells

To examine whether LLC-Exo can stimulate MDSC expansion in BM cells, we cultured LLC-Exo and 3T3-Exo with mouse BM cells in an inducing medium for seven days and the percentage of MDSC (CD11b^+^Gr-1^+^) in BM cells was determined by flow cytometry analysis. Furthermore, TEX from murine B lymphoma cell A20, murine melanoma cell B16, murine colorectal cancer cell CT26 (Supplementary Fig. 1A, B) were cultured with BM cells and MDSC percentages were characterised. We found there are more CD11B^+^Gr-1^+^ cells in BM cells cultured with LLC-Exo when compared to BM cells cultured with 3T3-Exo (Fig. 2A). In addition, BM cells cultured with TEXs from murine colorectal cancer cells (CT26-Exo) promote MDSC expansion too (Fig. 2A). We then evaluated whether these CD11B^+^Gr-1^+^ cells are functional to inhibit CD8^+^ T cell activation. CD11B^+^Gr-1^+^ MDSC were sorted out and cultured with mouse CD8^+^ T cells activated by anti-CD3 and anti-CD28 antibodies for another 48-hour. The proliferation and IFN-γ production of CD8^+^ T cells were detected to indicate the state of activation. The result showed that the proliferation of CD8^+^ T cells was almost stagnated by MDSC cells incubated in the presence of LLC-Exo (Fig. B). And the IFN-γ production of CD8^+^ T cells was evidently inhibited by MDSC cells incubated in the presence of LLC-Exo (Fig. 2C). These data indicated that TEX from LLC cells can induce functional MDSC expansion in mice BM cells.

**Figure 2.**
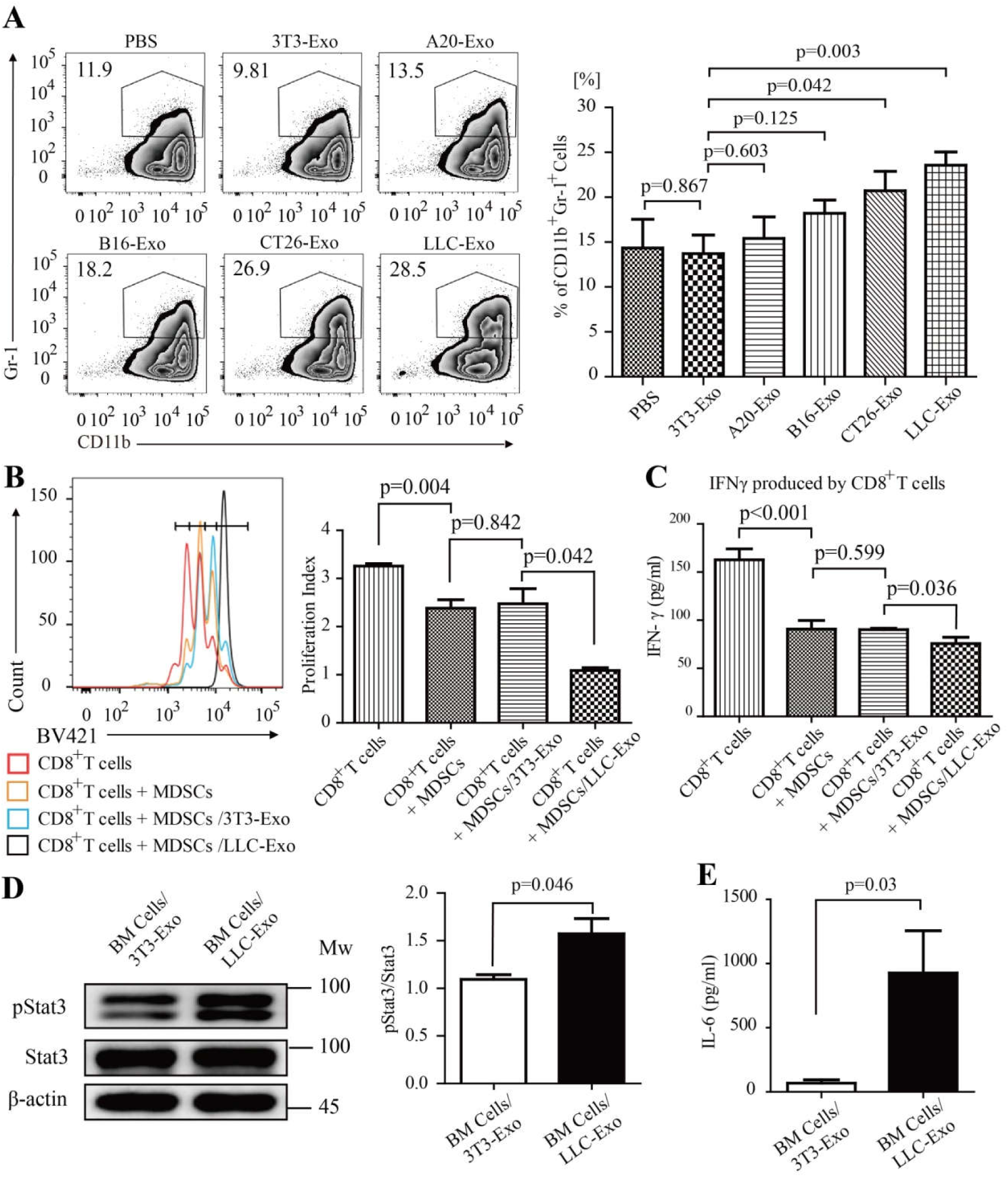
LLC-Exo promote MDSC expansion through activating the IL-6/Stat3 pathway. Mice bone marrow cells (5×10^5^ cells / mL) were cultured in an MDSC inducing RPMI 1640 medium (10% exosome-free FBS, 20 ng/ml GM-CSF) with exosomes (50 μg/mL) purified from 3T3 cells or other four tumour cell lines for 7 days. (**A**) The percentage of MDSC (CD11b^+^Gr-1^+^) was then determined using FACS. (**B**) CD11B^+^Gr-1^+^ MDSC were sorted using FACS and cultured with activated CD8^+^ T cells (3×10^4^ CD8^+^ T cells, 1×10^4^ MDSC cells, 200 μL/well) for 48-hours. The proliferation of CD8^+^ T cell was measured by labelled with CTV dye. (**C**) The concentration of IFN-γ in the supernatant was determined by ELISA. (**D**) After seven days in culture, the Y705-pStat3 protein in bone marrow cells was determined by using Western blotting. (**E**) The supernatants of seven-day cultures were collected and IL-6 concentration produced by bone marrow cells was determined by ELISA. A two-tailed student t test was used and all experiments in this figure were repeated for three times. Error bars represent the mean ± sem.

We then considered the molecular mechanism through which LLC-Exo stimulate MDSC expansion in BM cells. STAT3 has been proposed to be the main regulator of MDSC expansion through stimulation of myelopoiesis and inhibition of the differentiation of mature myeloid cells(*Guo et al., 2018a; Kortylewski et al., 2005; L. Li et al., 2014; Liu et al., 2010; Nefedova et al., 2005*). Tyrosine705 phosphorylation in STAT3 (pSTAT3 Y705) were shown to induce MDSC expansion(*Nefedova et al., 2007*). Consequently, we collected whole lysates of mice BM cells cultured with LLC-Exo or 3T3-Exo for seven days and the STAT3 with phosphorylated Y705 was determined with Western blotting. We found the amount of Y705 phosphorylated STAT3 to be significantly up-regulated in BM cells cultured with LLC-Exo (Fig. 2D). This indicated that LLC-Exo promote MDSC expansion through STAT3 pathway activation in mice BM cells.

It was then necessary to question what mechanism mediated LLC-Exo to enhance STAT3 activation in BM cells. Studies have demonstrated that various factors including GM-CSF(*Serafini et al., 2004*), VEGF(*D. Gabrilovich et al., 1998*), IL-10(*L. Wu et al., 2017*) and IL-6(*Menetrier-Caux et al., 1998; L. Wu et al., 2017*) produced mainly by tumour cells will induce MDSC expansion through JAK2-STAT3 pathway activation(*Kortylewski et al., 2005; Nefedova et al., 2005*). To that end, we detected concentration of these factors in the supernatant of BM cells cultured with LLC-Exo or 3T3-Exo by ELISA. Given that we had already added GM-CSF into the culture medium of BM cells, we did not check the concentration of GM-CSF. The findings were that there are evidently higher amount of IL-6 in medium of BM cells cultured with LLC-Exo (Fig. 2E) and little VEGF, IL-10 in mediums of BM cells cultured with LLC-Exo and 3T3-Exo (Supplementary Fig. 2D, E). To verify the original source of IL-6, we detected IL-6 concentration in LLC-Exo and 3T3-Exo by ELISA and found almost no IL-6 producing in the two exosomes (Supplementary Fig. 2F). The results convinced us that IL-6 is autocrine by the mice BM cells. The conclusion was that LLC-Exo phagocytosed by BM cells stimulates these cells to produce more IL-6 which probably triggers JAK2-STAT3 pathway into activating more vividly and finally promotes more MDSC expansion in mice BM cells.

### LLC-Exos carry a specific micoRNA signature containing high amount of miRNA-21a molecule

The next stage was to identify a mechanism that LLC-Exo induce mice BM cells to produce more IL-6. In 2009, Xiang et al. reported that TEX from mice mammary carcinoma cells can induce IL-6 production and MDSC accumulation depending on TGF-beta molecules in TEX(*Xiang et al., 2009*). In 2010, Chalmin et al. found that TEX associated heat shock protein 72 (Hsp72) triggers STAT3 activation in MDSC in a TLR2/MyD88-dependent manner through autocrine production of IL-6(*Chalmin et al., 2010*). We then checked the expression of molecules including Hsp72, Hsp90 and TGF-β in TEXs by Western blotting and ELISA accordingly and found that the amount of these molecules in LLC-Exo and 3T3-Exo were similar (Supplementary Fig.3A, B). This indicated that there could be an unknown mechanism mediating LLC-Exo function to regulate IL-6 production in BM cells. Recent studies pointed out miRNA are potential biological functional molecules in exosomes(*Valadi et al., 2007*) and also play an important roles to regulate MDSC expansion and activation(*Guo et al., 2018a; Guo et al., 2018b; L. Li et al., 2014*). Therefore, we are curious about the possibility that TEX associated miRNA might play a role in the regulation of IL-6 production in mice BM cells. Since our previous result showed that LLC-Exo and CT26-Exo both can stimulate MDSC expansion significantly, we sequenced miRNA expressing in both TEX from LLC and CT26 cells produced by RNA-Seq technology (GZRB2016081201901SSQ, RiboBio in Guangzhou, China). We observed fourteen miRNA including miR-21a molecules to be highly expressing in both TEX from LLC and CT26 cells (Fig. 3A). miR-21a plays a main role in cell proliferation, migration, invasion and apoptosis and has been up-regulated in many cancer cells(*Selcuklu et al., 2009; Si et al., 2007*). Furthermore, studies reported that miR-21a induces the expansion both of monocytic and granulocytic MDSC via targeting phosphatase and tensin homolog (PTEN), leading to STAT3 activation(*L. Li et al., 2014*). Consequently, we further validated miR-21a expression in LLC-Exo and other TEX through quantitative real-time PCR (qPCR) analysis. When compared to the amount of miR-21a expressed in 3T3-Exo and other TEXs, there were distinctly higher amounts of miR-21a contained in LLC-Exo (Fig. 3B).The content of miR-21a in cell cytoplasm was, however, totally different from what was found in cell exosomes (Supplementary Fig. 3C). We then observed that the content of miR-21a in mice BM cells culturing in an MDSC inducing medium in vitro for seven days was dynamically increasing (Supplementary Fig. 3D). Combining the above results, we considered that the miR-21a contained in LLC-Exo might have a potential role in regulating MDSC expansion.

**Figure 3.**
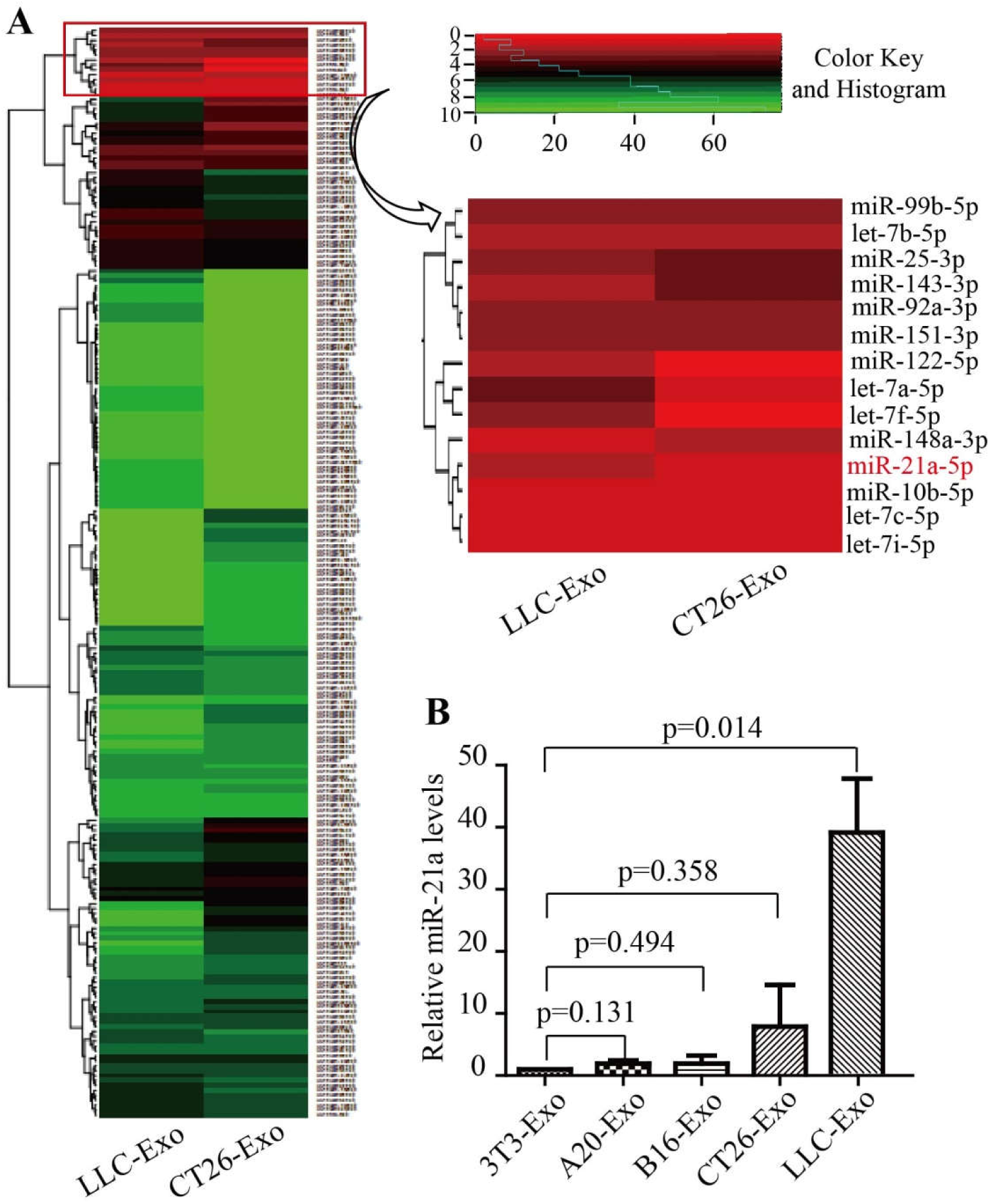
The relative amount of miRNA-21a in LLC-Exo is high. (**A**) Total RNA extracted from either LLC-Exo or CT26-Exo was used for microRNA-Seq assay. A heat map illustrated the cluster of miRNAs increasing expression both in LLC-Exo and CT26-Exo. (**B**) The expression of miR-21a in exosomes from 3T3 cells and other different cell lines was verified by quantitative RT-PCR. A two-tailed student t test was used and the data is representative of three independent experiments. Error bars represent the mean ± sem.

### LLC-Exo-associated miR-21a promote MDSC expansion via stimulating IL-6 production in mice BM cells

To validate the effect of miR-21a on regulating MDSC expansion, we cultured mice BM cells with an MDSC inducing medium containing miR-21a mimics (100nM) (miR-21a mimic) or the control scrambled oligonucleotides (Cont mimic) for seven days. The percentage of MDSC was determined using FACS analysis. Results showed that BM cells with miR-21a over-expression by miRNA mimics transfection could promote MDSC expansion (Fig. 4A). Furthermore, there were more phosphorylated Y705 STAT3 and IL-6 production in BM cells with miR-21a over-expression (Fig. 4B, C). We then successfully deleted miR-21a encoding gene in LLC cells genome (LLC-21a-KO) by CRISPR/Cas9 system (Supplementary Fig. 4A). The expression of miR-21a in exosomes from LLC-21a-KO cells (LLC-21a-KO-Exo) had almost vanished when compared to that in LLC-Exo (Supplementary Fig. 4B). We cultured mice BM cells in a MDSC induction medium with equal amounts of LLC-Exo and LLC-21a-KO-Exo respectively for seven days and the percentage of MDSCs was determined by FACS analysis. We observed that the percentage of CD11B^+^Gr-1^+^ population in BM cells with LLC-21a-KO-Exo was remarkably reduced when compared to that in BM cells cultured with LLC-Exo (Fig.4D). What is even more notable is that phosphorylated Y705 STAT3 and IL-6 production in BM cells cultured with LLC-21a-KO-Exo was also sharply reduced (Fig. 4E, F). These results demonstrated that miR-21a molecules containing LLC-Exo phagocytosed by mice BM cells can promote MDSC expansion through STAT3 activations which is triggered by autocrine IL-6.

**Figure 4.**
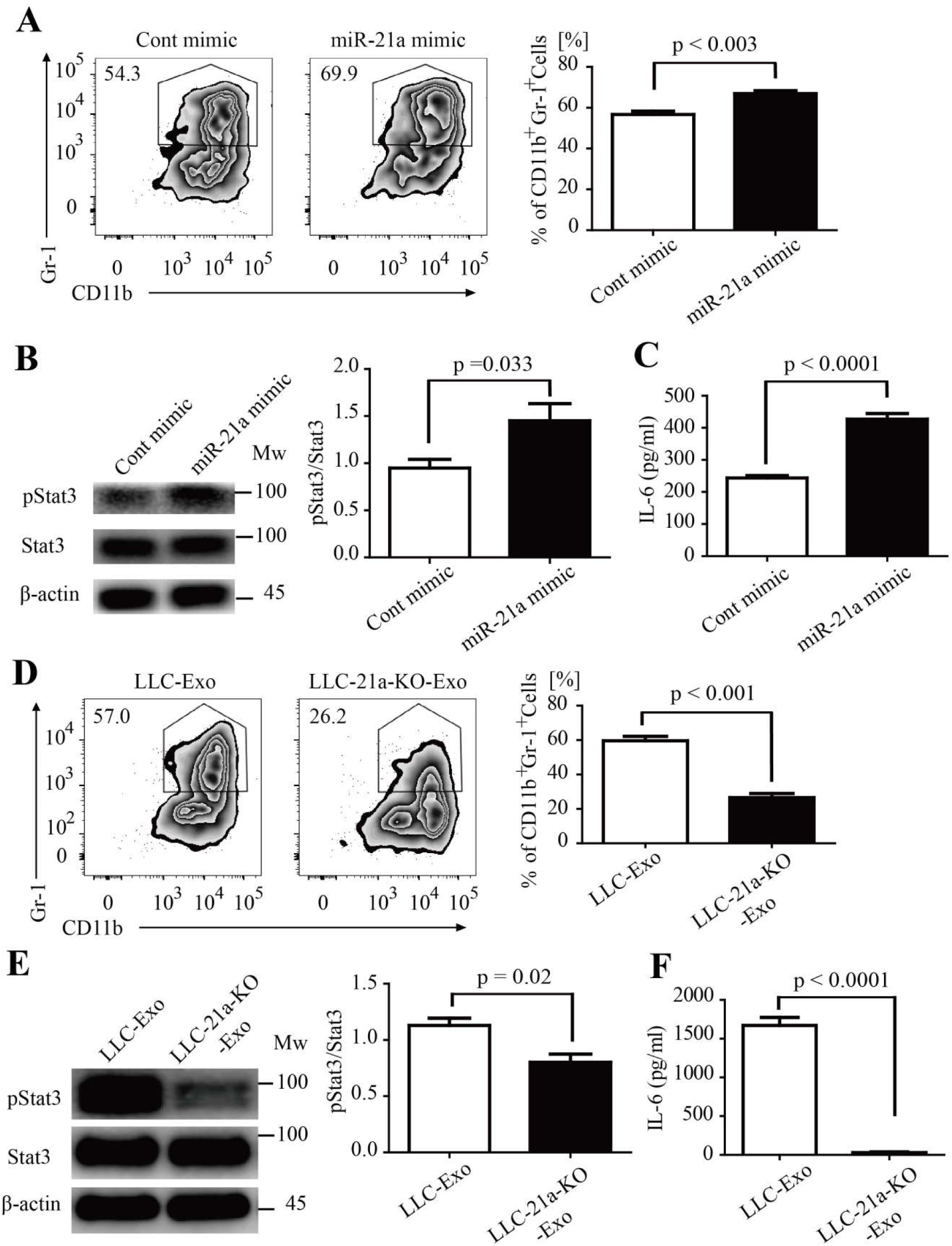
miRNA-21a from LLC-Exo enhances the MDSC accumulation through activating the IL-6/Stat3 pathway. Mice bone marrow cells (5×10^5^ cells/mL) were cultured in an MDSC inducing RPMI 1640 medium containing 10% exosome-free FBS, GM-CSF (20 ng/mL). Mice bone marrow cells were transfected with 100 nM miR-21a mimic or control scrambled oligonucleotides (Cont mimic). After seven days, (**A**) CD11b^+^Gr-1^+^ MDSC were evaluated using FACS. (**B**) Expression of STAT3 and pSTAT3 in the induced BM cells was detected by Western blotting. (**C**) IL-6 concentration in the supernatant of the medium was determined by ELISA. We constructed miR-21a-knockout LLC cell clones (LLC-21a-KO) using the CRISPR/Cas9 genome editing system. Mice bone marrow cells were cultured in an MDSC inducing medium with exosomes derived from normal LLC cells or LLC-21a-KO cells (LLC-21a-KO-Exo) for seven days as described above. (**D**) The percentage of CD11b^+^Gr-1^+^ MDSCs was determined using FACS. (**E**) The phosphorylation of Y705 in STAT3 protein in bone marrow cells was determined using Western blotting. (**F**) IL-6 concentration in the supernatant MDSC medium was determined by ELISA. A two-tailed student t test was used and all experiments in this figure were repeated for three times. Error bars represent the mean ± sem.

### PDCD4 is targeted by miR-21a in mice BM cells

The miRNA are small noncoding RNA, 19-24nt in length, with canonical binding to their target messenger RNA to inhibit gene-expression(*Bartel, 2009*). We were curious about molecules which are targeted by miR-21a to induce IL-6 production in mice BM cells. Previous studies indicate that miR-21a, as potential regulator of MDSC expansion and immunosuppressive function(*El Gazzar, 2014*), was proved to have an effect on cells via targeting PTEN molecules(*Guo et al., 2018a; L. Li et al., 2014*). In our study, however, we did not observe any significant difference of PTEN expression in the BM cells cultured with LLC-Exo compared to BM cells cultured with 3T3-Exo (Supplementary Fig. 5A). Recent investigations have found that miR-21a plays a key role in macrophages to regulate anti-inflammation response via targeting PDCD4 (*Das et al., 2014; Merline et al., 2011; Sheedy et al., 2010*). In glioblastoma cells increased PDCD4 expression led to the attenuation of activator protein-1 (AP-1) transcription by inhibiting c-Jun phosphorylation and resulted in a concomitant decrease in the expression of AP-1-target genes like IL-6(*Dikshit et al., 2013; Wang et al., 2012*). We also observed that the amount of PDCD4 was reduced in BM cells cultured with LLC-Exo which contain higher amount of miR-21a molecules compared to BM cells cultured with 3T3-Exo (Fig. 5A). PDCD4 recovered its expression in BM cells when they were cultured with LLC-21a-KO-Exo in which miR-21a had vanished (Fig. 5A). Furthermore, we found there were conserved miR-21a binding site existing in 3’UTR of mouse PDCD4 mRNA (Supplementary Fig. 5B). To further verify the binding of miR-21a to 3’UTR of PDCD4 mRNA, we constructed a reporting system with PDCD4 3’-UTR containing the sequences predicted as miR-21a binding site inserted into the downstream of firefly luciferase encoding gene (PDCD4-WT-UTR). A plasmid with a mutant in the sequence of miR-21a binding site inserted into the downstream of firefly gene (PDCD4-MT-UTR) that was used as the control (Supplementary Fig. 5B). The relative activity of firefly luciferase was normalising to the activity of renilla luciferase in 293T cells transfected with luciferase reporter system and miR-21a mimics simultaneously. We observed that firefly activity in 293T cells transfected with miR-21a mimics and a reporting system with PDCD4-WT-UTR (miR-21a mimic + PDCD4-WT-UTR) was evidently inhibited when compared to that in cells transfected with scramble oligonucleotides (Cont mimic + PDCD4-WT-UTR) (Fig. 5B). The inhibition disappeared when cells are transfected with a reporting system containing a mutant sequence of PDCD4 3’-UTR and with miR-21a mimics simultaneously (miR-21a mimic + PDCD4-MT-UTR) (Fig. 5b).This indicated that the binding of miR-21a to a conserved binding site of PDCD4 located in firefly mRNA 3’-UTR can inhibit its expression. To characterise the role of miR-21a plays in regulating PDCD4 expression, we added miR-21a mimics (100 nM) and scramble oligonucleotides to 3T3 cells respectively (3T3-miR-21a mimic or 3T3-Cont mimic) for three days. The amounts of PDCD4 mRNA and protein molecules were then determined by relative quantitative RT-PCR and Western blotting analysis. We found that PDCD4 protein in 3T3 cells with over-dosed miR-21a (3T3-miR-21a mimic) was heavily reduced but PDCD4 mRNA molecules were equal in number when compared to those in 3T3 cells transfected with scramble oligonucleotides (3T3-Cont mimic) (Fig. 5C, D). We also checked the amount of PDCD4 mRNA and protein molecules in wild type LLC and LLC with deletion of miR-21a (LLC-21a-KO). Elimination of miR-21a in LLC caused the PDCD4 protein amount to increase to an obviously higher level when compared to that in wild type LLC (Fig. 5F). The PDCD4 mRNA amounts in wild type LLC and LLC-21a-KO showed no difference (Fig. 5E). These results indicate that miR-21a inhibits PDCD4 expression post-transcriptionally.

**Figure 5.**
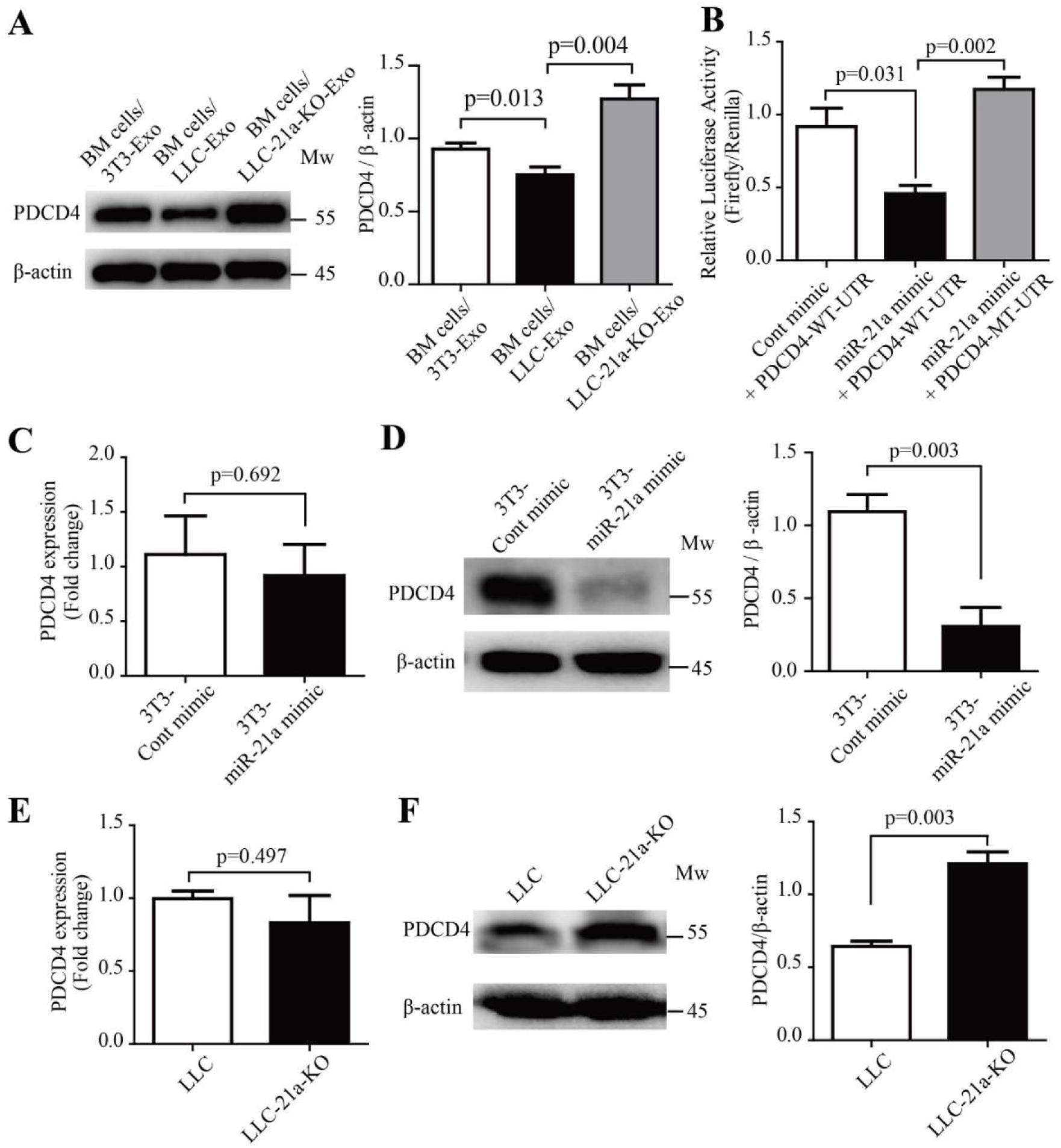
miR-21a down-regulates PDCD4 protein via binding to the 3’-UTR of PDCD4 mRNA during MDSC expansion. (**A**) Mice bone marrow cells (5×10^5^ cells/mL) were cultured in an MDSC inducing RPMI 1640 medium with exosomes (50 μg/mL) of 3T3-Exo, LLC-Exo or LLC-21a-KO-Exo for seven days. The relative abundance of PDCD4 protein in induced BM cells was then determined by Western blotting. (**B**) The entire 3′-UTR of murine PDCD4 (PDCD4-WT-UTR) or 3′-UTR containing mutations in the miR-21a seed region-binding sites (PDCD4-MT-UTR) were cloned into the downstream of the firefly luciferase reporter gene in pEZX-MT06 vector. Plasmids (500 ng) and synthetic miR-21a mimic RNA fragments (miR-21a mimic) or control RNA fragments (Cont mimic) (100 nM) were transiently transfected into 293T cells (1×10^5^ cells/mL). The luciferase activity in 293T cells was measured 72 hours later. (**C-D**) The 3T3 cells (5×10^5^ cells/mL) were transiently transfected with 100 nM miR-21a mimic or Cont mimic. Forty-eight hours later, the 3T3 cells were harvested and subjected to quantitative PCR analysis and Western blotting to determine the relative abundance of mRNA and protein molecules of PDCD4 respectively. (**E-F**) Quantitative PCR analysis and Western blotting were used to determine the relative abundances of mRNA and protein molecules of PDCD4 in LLC cells and LLC-21a-KO cells respectively. A two-tailed student t test was used and all experiments in this figure were repeated for three times. Error bars represent the mean ± sem.

### PDCD4 inhibits MDSC expansion in mice BM cells via down-regulating IL-6 production

PDCD4 is well known as a tumour suppressor and potential target of anticancer therapies(*Lankat-Buttgereit and Goke, 2009; Wang and Yang, 2018*) and also responds to inflammatory stimuli (*Cohen and Prince, 2013; van den Bosch et al., 2014*). In glioblastoma cells, PDCD4 inhibits IL-6 production via attenuation of AP-1 transcription(*Dikshit et al., 2013*). The function of PDCD4 to regulate MDSCs expansion and IL-6 production in mice BM cells is, however, inscient. To determine the function of PDCD4 in mice BM cells, we first constructed a lentivirus vector which delivered shRNA segments against mouse PDCD4 (shPDCD4). PDCD4 expression was down-regulated in cells with the infection of shPDCD4 lentivirus (Supplementary Fig. 6A, B). Fresh mouse BM cells were infected with shPDCD4 lentivirus or control shNC virus particles, and then cultured in an MDSC inducing medium. Seven days later, the percentage of CD11B^+^Gr-1^+^ MDSC population was determined by FACS analysis. The amount of phosphorylated Y705 STAT3 molecules in these induced BM cells was determined using Western blotting, and IL-6 production was determined using ELISA. The results showed that PDCD4 reduction led to heavier expansion of MDSC in BM cells with an infection of shPDCD4 lentivirus (Fig. 6A). IL-6 production and the amount of phosphorylated Y705 STAT3 had clearly increased in the mice BM cells with PDCD4 reduction (Fig. 6B, C). Furthermore, we constructed a lentivirus vector (LV-PDCD4) to induce PDCD4 overexpression in mice BM cells (Fig. 6E). Fresh mouse BM cells were infected with LV-PDCD4 or control LV-NC virus particles, and then cultured in an MDSC inducing medium. Then the percentage of MDSC population, amount of phosphorylated Y705 STAT3 molecules and IL-6 production were checked. We found that overexpression of PDCD4 in BM cells evidently suppresses the expansion of MDSC (Fig. 6D) and that IL-6 production and phosphorylated Y705 STAT3 molecules are accordingly reduced in these cells (Fig. 6E, F). These results indicated that PDCD4 is a critical inhibitor to negatively regulate expansion of MDSC via down-regulating activation of IL-6/STAT3 pathway in mice BM cells. We still did not, however, confirm the mechanism of LLC-Exo to promote MDSC expansion whether or not through PDCD4 reduction in mice BM cells. To confirm this question, we employed a LV-NC or LV-PDCD4 virus particle to infect mice BM cells which were cultured in an MDSC inducing medium with a mixture of LLC-Exo for seven days. Then the percentage of MDSC population, amount of phosphorylated Y705 STAT3 molecules and IL-6 production in these cells was determined. We found that MDSC expansion was blunted in BM cells with overdose exogenous PDCD4 (Fig. 6G) and that the amount of phosphorylated Y705 STAT3 and IL-6 production both decreased in these cells (Fig. 6H, I). Since previous studies reported that in glioblastoma cells PDCD4 could slash AP-1 transcription activity by inhibiting c-Jun phosphorylation and result in reduced IL-6 expression(*Dikshit et al., 2013*), we proposed that PDCD4 reducing in BM cells releases a brake on AP-1 transcription by up-regulating c-Jun phosphorylation and results in IL-6 autocrine production promotion and finally stimulates more STAT3 activation in these cells.

**Figure 6.**
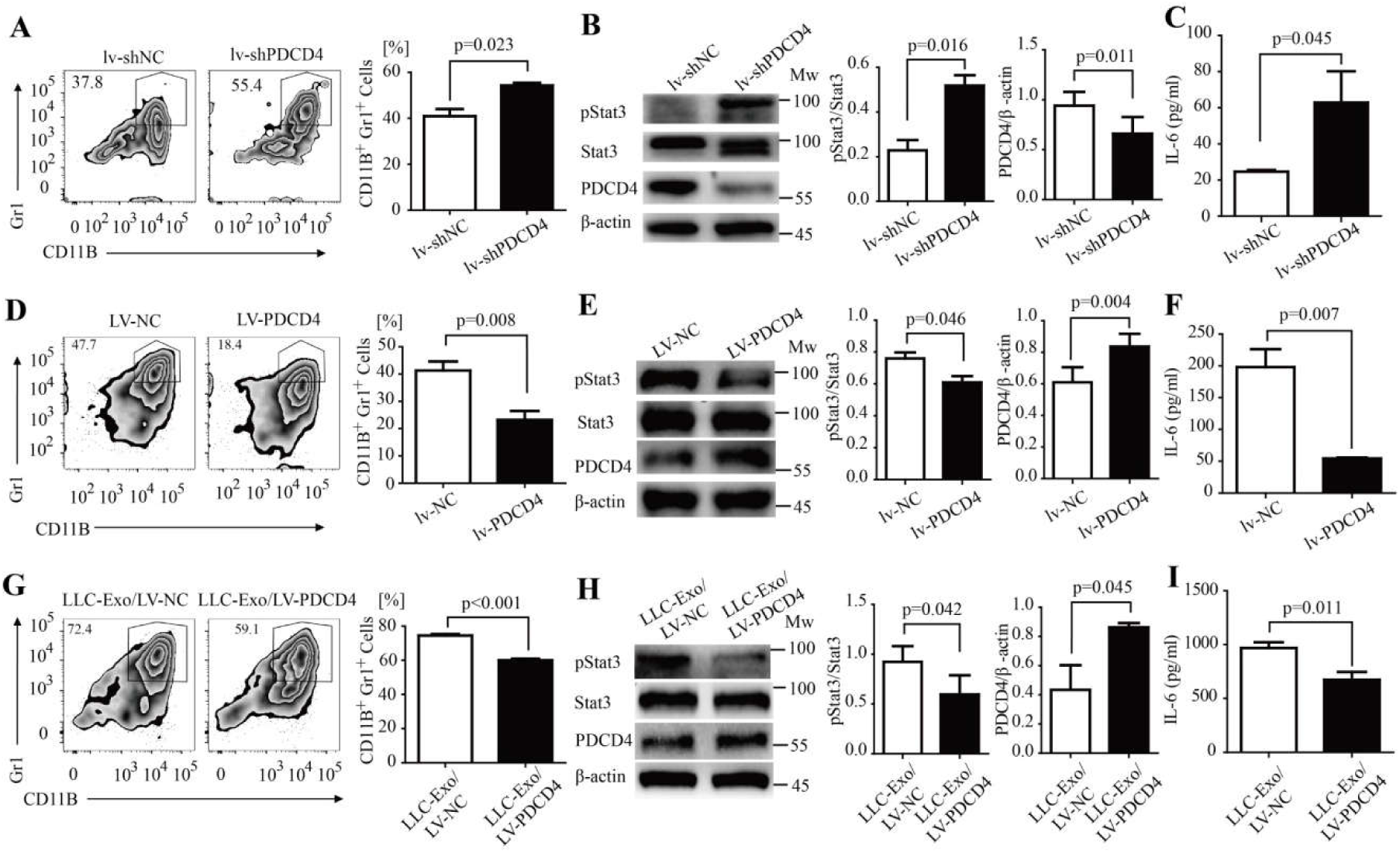
PDCD4 negatively regulated MDSC expansion through activation of the IL-6/Stat3 pathway. (**A-C**) A designed shRNA sequence to inhibit PDCD4 transcription (shPDCD4) and control sequence (shNC) was cloned into the MISSION® TRC2 pLKO.5-puro empty vector (SHC201, Sigma). The lentivirus was packaged with these vectors. Mice BM cells were isolated and infected with shPDCD4 or shNC lentivirus particles (lv-shPDCD4 or lv-shNC). These cells (5×10^5^ cells/mL) were then cultured in the MDSC inducing medium for seven days. The percentage of CD11b^+^Gr-1^+^ MDSCs was determined using FACS analysis (**A**).The amount of Y705-pStat3 protein in induced BM cells was determined by Western blotting (**B**). IL-6 concentration in the supernatant was determined using ELISA (**C**). (**D-F**) The whole sequence encoding for PDCD4 was cloned into a lentivirus vector FtetUGW-T (lv-PDCD4).The lentivirus was packaged with a lv-PDCD4 vector or mock vector as control (lv-NC). Mice BM cells were isolated and infected with lv-PDCD4 or lv-NC lentivirus particles. These cells (5×10^5^ cells/mL) were cultured in the MDSC inducing medium for seven days. The percentage of CD11b^+^Gr-1^+^ MDSCs was determined using FACS analysis (**D**).The amount of Y705-pStat3 protein in induced BM cells was determined using Western blotting (**E**). IL-6 concentration in the supernatant was determined using ELISA (**F**). (**G-I**) Mouse BM cells were isolated and then infected with lv-PDCD4 or lv-NC lentivirus particles. Cells (5×10^5^ cells/mL) were cultured in MDSC inducing medium with LLC-Exo (50 μg/mL) for seven days. The percentage of CD11b^+^Gr-1^+^ MDSCs was determined using FACS analysis (**G**). The Y705-pStat3 protein in induced BM cells was determined using Western blotting (**H**). IL-6 concentration in the supernatant was determined using ELISA (**I**). A two-tailed student t test was used and all experiments in this figure were repeated for three times. Error bars represent the mean ± sem.

### TDX from human lung cancer cells induce human monocyte differentiation into hMDSC

We extracted exosomes from several different human lung cancer cell lines including lung carcinoma 95D, lung mucoepidermoid carcinoma H292 and bronchioalveolar carcinoma H358 cells and extracted exosomes from human normal embryonic kidney epithelial cells 293T as the control (95D-Exo, H292-Exo, H358-Exo, 293T-Exo, Supplementary Fig. 7A, B). Human CD14^+^ monocytes purified from the peripheral blood of healthy donors were cultured with an equal quantity of above exosomes respectively in an MDSC inducing medium with hGM-CSF and hIL-4. Four days later, the percentage of human MDSC population (CD11b^+^CD33^+^HLA-DR^−^) was assessed by use of FACS. The results showed that TDX from human lung cancer cells can induce human monocyte differentiation into hMDSC (Fig. 7A). Furthermore, we checked miR-21a expression in TDX and cancer cells by relative quantitive RT-PCR analysis and found the amount of miR-21a in TDX and cancer cells (95D-Exo, H292-Exo, H358-Exo) to be higher than those in 293T-Exo or normal 293T cells (Fig. 7B, Supplementary Fig. 7C). These results hinted that miR-21a might play a potential role in lung cancer exosomes and trigger hMDSC expansion.

**Figure 7.**
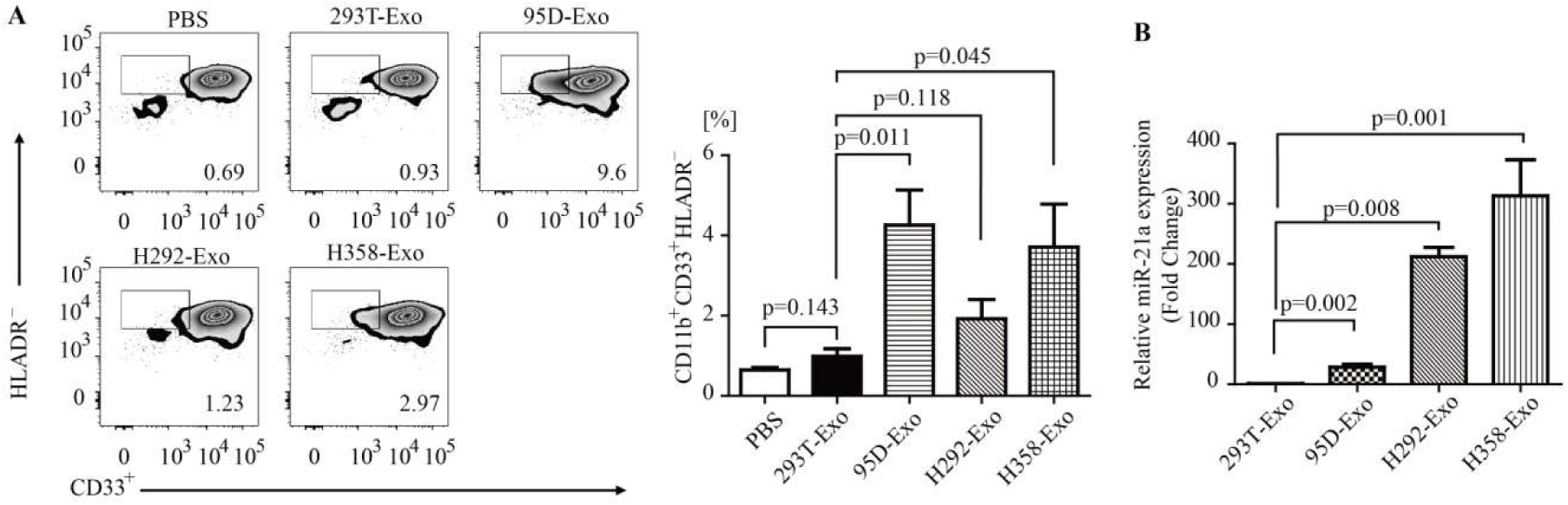
Exosomes derived from human lung cancer cell lines triggered human MDSC expansion. (**A-B**) PBMCs from normal human donors were cultured (5×10^5^ cells/mL) in the PRMI1640 medium with 10% exosome-free FBS, hIL-4 (20 ng/mL), hGM-CSF (50 ng/mL) and exosomes (50 μg/mL) purified from 293T cells, 95D cells, H292 cells or H358 cells for 4 days. **A** The percentage of hMDSC (CD11b^+^CD33^+^HLADR^-^) was determined using FACS. **B** The expression of miR-21a in exosomes from 293T cells, 95D cells, H292 cells and H358 cells was verified by quantitative RT-PCR. A two-tailed student t test was used and data is representative of three independent experiments. Error bars represent the mean ± sem.

## Discussion

As a well-known mediator of intercellular communication, tumour-derived exosomes (TEX) are crucial in multiple aspects of tumour progression and especially in regulating tumour associated immune response(*Hingorani, 2015; Steinbichler et al., 2017*). Numerous studies have indicated that TEX play a dual role in modulating tumour immunity(*Liu et al., 2015*). On one hand, TEX can stimulate tumour-specific CTL responses by delivering native tumour-associated proteinsto dentritic cells(*Andre et al., 2002*). On the other, more and more recent studies indicate that TEX is a novel element to mediate immune-inhibitory activity in tumour cells(*Ashiru et al., 2010; Muller et al., 2016; Wieckowski et al., 2009; Zhou et al., 2014*). TEX are able to skew the differentiation of myeloid precursor cells into MDSC (*Chalmin et al., 2010; Guo et al., 2018a; Guo et al., 2018b; Liu et al., 2010; Xiang et al., 2009*), influence macrophages function(*Chow et al., 2014*), inhibit NK cell cytotoxicity(*Ashiru et al., 2010*), induce formation of regulatory T cells(*Wieckowski et al., 2009*) or regulatory B cells(*Y. Li et al., 2015*) and finally favour developing a tumour-mediated immune suppression environment.

MDSC are a heterogeneous population comprising of immature myeloid cells generated in the bone marrow with remarkable ability to suppress T-cell dependent anti-tumour response(*D. I. Gabrilovich and Nagaraj, 2009; Makarenkova et al., 2006; Ostrand-Rosenberg and Sinha, 2009*). In the study, MDSC was found to heavily accumulate in the peripheral blood of most patients with renal cell carcinoma(*Ochoa et al., 2007*), bladder carcinoma(*Yuan et al., 2011*), multiple myeloma(*Brimnes et al., 2010*), lung cancer(*de Goeje et al., 2015; Heuvers et al., 2013; Vetsika et al., 2014*) and other tumours(*Diaz-Montero et al., 2009*). TEX has been documented in response to MDSC expansion during the progress of the tumour^9-13^. Murine mammary adenocarcinoma cells promote MDSC accumulation through exosomal prostaglandin E2 (PGE2) and TGF-β molecules(*Xiang et al., 2009*). MiR-10a, miR-21a, miR-29a and miR-92a in TEX of glioblastoma are responsible for the expansion and immune suppressive function of MDSCs(*Guo et al., 2018a; Guo et al., 2018b*). In this study, we found that LLC TEX favours tumour growth probably through its ability to stimulate MDSC expansion in mice bone marrow then induce systemic accumulation of immune suppressive MDSCs in tumour-bearing mice (Fig.1 and Fig.2A-C).

The JAK-STAT3 is one of critical pathways to mediate MDSC expansion(*Kortylewski et al., 2005; Nefedova et al., 2005*). The Y705 phosphorylation of STAT3 in myeloid progenitor cells triggers its proliferation and survival through up-regulating several targeted gene expressions (cyclin D1, MYC, BCL-XL, survivin)(*D. I. Gabrilovich and Nagaraj, 2009; Imada and Leonard, 2000; Rane and Reddy, 2000*). Meanwhile, the JAK-STAT3 pathway activation skews the differentiation of the myeloid precursor into MDSC by inducing S100A8 and S100A9 expression (*Cheng et al., 2008; D. I. Gabrilovich and Nagaraj, 2009; Sinha et al., 2008*). Recent studies have reported that microRNAs in hypoxia-induced glioblastoma exosomes induce MDSC expansion via directly targeting PTEN and Prkar1a to lead to the STAT3 pathway activating(*Guo et al., 2018a; Guo et al., 2018b*). In our study, we found that LLC-Exo enhanced MDSC expansion through stimulating more STAT3Y705 phosphorylation in mice BM cells (Fig. 2D). A variety of factors converged on JAK-STAT3 activation to induce MDSC expansion including COX2(*Rodriguez et al., 2005*), prostaglandins(*Sinha et al., 2007*), SCF(*Pan et al., 2008*), M-CSF(*Menetrier-Caux et al., 1998*), IL-6(*Menetrier-Caux et al., 1998; C. T. Wu et al., 2012; L. Wu et al., 2017*), GM-CSF(*Serafini et al., 2004*) and VEGF(*D. Gabrilovich et al., 1998*). We found that mice BM cells treated with LLC-Exo could produce more IL-6 and lead to more Y705 STAT3 phosphorylation in these cells (Fig.2 D, E). We consider, therefore, that autocrine IL-6 of BM cells could trigger the JAK2-STAT3 pathway activating in myeloid progenitor cells and then inducing MDSC expansion.

It is important to consider what mechanism in LLC-Exo stimulates IL-6 production in mice BM cells. The results shown here demonstrate that miR-21a, which is collected in LLC-Exo, has a critical role to regulate IL-6 production in mice BM cells (Fig. 4). Overexpression of miR-21a in mice BM cells with microRNA mimics induces up-regulation of IL-6 production in these cells (Fig. 4C). Otherwise, without mature miR-21a in LLC-Exo, the production of IL-6 in mice BM cells treated by TEX almost vanished (Fig. 4F). MiR-21a is an abundantly expressed miRNA in mammalian cells, which has been shown to be the most commonly upregulated miRNA in solid and haematological malignancies(*Selcuklu et al., 2009; Si et al., 2007*). In our study, we also find it is abundant in several different tumour cell lines (Supplementary 3c, 7c). It is well documented that miR-21a is an oncogenic miRNA to regulate cancer cell proliferation, migration and apoptosis by suppressing the expression of tumour suppressors including PTEN(*Mao et al., 2017*) and PDCD4(*Asangani et al., 2008; Mao et al., 2017*). Recently miR-21a emerges to have a role in regulating inflammatory response and to stimulate production of anti-inflammatory cytokine IL-10 in macrophage through miR-21/PDCD4 axis(*Das et al., 2014*). Furthermore, miR-21 shows a synergistic effect with miR-155 to induce MDSC differentiation by targeting SHIP-1 and PTEN, which leads to STAT3 activation(*L. Li et al., 2014*). In this study, we confirm that miR-21a stimulates mice BM cells to produce IL-6 via targeting PDCD4 (Fig. 5).

PDCD4 was for several years mainly known as a tumour suppressor gene and potential target for anticancer therapies (*Lankat-Buttgereit and Goke, 2009; Wang and Yang, 2018*). Recent reports show that PDCD4 is response to inflammatory stimuli and can regulate inflammation(*Cohen and Prince, 2013; van den Bosch et al., 2014*). In this study, we validate that PDCD4 is indeed a brake on MDSC expansion via IL-6 production reduction leading to STAT3 phosphorylation inhibiting in BM cells (Fig. 6 D-F). PDCD4 reduction in BM cells leads to heavier expansion of MDSC with an obvious increase of IL-6 production and Y705 STAT3 phosphorylation (Fig. 6A-C). PDCD4 is a well-known AP-1 inhibitor(*Talotta et al., 2009; Zhang et al., 2013*), and transcription of IL-6 gene is under the control of AP-1(*Dikshit et al., 2013*). In addition, miR-21a in macrophages silences PDCD4, favouring c-Jun-AP1 activity, which in turn results in elevated production of anti-inflammatory cytokine IL-10(*Das et al., 2014*). As a result, we concluded that miR-21a in LLC-Exo mediates MDSC expansion via down-regulating PDCD4 to promote IL-6 production in mice BM cells.

## Materials and methods

### Mice

Female C57BL/6 mice between six to eight weeks old were purchased from Daping Hospital. Mice were maintained in the Army Medical University under specific-pathogen-free (SPF) conditions. All animal experiments were conducted according to protocols approved by the Medicine Animal Care Committee of the Army Medical University (Chongqing, China).

### Cell Culture

The mouse Lewis lung carcinoma LLC cells, colon carcinoma CT26 cells, B lymphoma A20 cells, melanoma B16 cells, embryo fibroblast NIH/3T3 cells, LLC-luc (an orthotopic luciferase stably expressed LLC) were obtained from the laboratory of Professor Wan of the Army Medical University of China. The human lung cancer 95D (high invasive ability) cells, lung mucoepidermoid carcinoma H292 cells, lung adenocarcinom H358 cells, embryonic kidney 293T cells were purchased from the Chinese Academy of Sciences Cell Bank. A20 cells were grown in RPMI 1640 medium (whereas all other cells were grown in DMEM) supplemented with 10% (v/v) FBS (Lonza) and with Pen/Strep Amphotericin B (Lonza) in an atmosphere of 95% air and 5% CO2 at 37°C.

### Exosome Isolation and Procedures

Cells were cultured in DMEM/1640 supplemented with 10% exosome-depleted FBS (EXO-FBS-250A-1, SBI System Biosciences, USA) using 10×10 cm^2^ disks (1×10^6^ cells per disk) for 48 hours. Differential centrifugation was then performed to isolate exosomes from supernatants of conditioned medium. Initial spins was centrifuged (4°C) at 600 g for ten minutes and then at 12000g for thirty minutes. The media were subsequently filtered using a 0.22-μm pore filter (Steriflip, Millipore). The collected media were then ultracentrifuged at 110,000 g for 2 h at 4 °C. The exosomes pellet was washed once in PBS, followed by a second step of ultracentrifugation at 110,000 g for 2 h at 4 °C. The final pellet contained the exosomes, which were re-suspended in PBS (sterile). The protein concentration of the exosomes was measured by Lowry protein assay (Bio-Rad). This protocol is mainly based on previous exosome isolation methods(*Cooks et al., 2018; Thery et al., 2006*).

### Electron Microscopy

Exosome preparations were loaded onto a carbon-coated electron microscopy grid and stained with uranyl-acetate solution (4%) for ten minutes. On the next day, the grids were observed with a transmission electron microscope Philips-FEI TecnaiT10 TEM.

### Exosome Labelling and Examination

Exosomes were labelled using the green lipophilic fluorescent dye PKH67 (Sigma) for five minutes and the reaction was stopped by the addition of exosome-depleted FBS. Bone marrow (BM) cells were incubated with the labelled exosomes for five hours, washed with PBS, fixed with 4% paraformaldehyde for twenty minutes and then stained with ProLongH Gold Antifade Reagent with a DAPI nuclear stain (Life Technologies, Grand Island, NY). Pictures were taken using a TCS-SP5 (Leica).

C57BL/6 mice of 6-8 weeks of age were injected intravenously with PKH67-labeled exosomes (100 μg/100 μL/mouse) or an equal volume of PBS. The mice were sacrificed twenty-four hours after injection. Cells from the spleen and bone marrow were recovered and subjected to FACS analysis. The percentage of cells containing exosomes was determined by counting FITC-positive cells.

### In-vivo Induction of MDSC by Exosomes

LLC-luc cells (1×10^6^ cells/100 μL/mouse) were co-injected with LLC-Exo or 3T3-Exo (100 μg/100 μL/mouse). On the fifteenth day post injection, cells from the spleen were recovered. The percentages of CD11b^+^Gr-1^+^ cells were examined using FACS analysis.

### In vivo Tumour Growth

The growth of tumour (LLC-luc cells) was monitored by two methods. The bioluminescence imaging of the tumour was carried out using the IVIS imaging system and calculated to total luminescent fluxes. The volume of tumour size was measured using callipers and calculated to total luminescent fluxes or according to the formula V = Length × Width^2^/2. Animals were sacrificed when the maximal allowable tumour size was reached for fifteen days.

### Generation of MDSC from Bone Marrow Progenitors

Bone marrow cells were obtained from the femurs and tibias of WT mice and cultured ex-vivo in 24-well plates (5×10^5^ cells/mL). BM cells culture medium was consisted of RPMI 1640 medium supplemented with 10% exosome-depleted FBS (EXO-FBS-250A-1, SBI), GM-CSF (20 ng/mL; peprotech) and with or without exosomes (50 μg/mL). Half of the volume of the medium was replaced by MDSC inducing medium every two days for a total if seven days in the culture.

### Generation of MDSC from Human Peripheral Blood Mononuclear Cell (PBMC)

The human MDSC were generated as previously described(*Spaggiari et al., 2009*). Briefly, PBMC were isolated from a source of healthy donors via density gradient centrifugation using Ficoll-Paque (tbdscience, China) according to the manufacturer’s protocol. CD14^+^ cells were positively selected from PBMC using anti-CD14-coated magnetic beads (Miltenyi Biotec, Bergisch Gladbach, Germany). Then cells were cultured in 24-well plates (5×10^5^ cells/mL). The culture medium consisted of RPMI 1640 supplemented with 10% exosome-depleted FBS, human GM-CSF (50 ng/mL; peprotech, Rocky Hill, NJ, USA), human IL-4 (20 ng/mL; peprotech) and with or without TEXs (50 μg/mL). After four days, the cultured cells were harvested and analysed by flow cytometry.

### Flow CytometryAnalysis of MDSC

For mouse MDSC, single-cell suspensions were stained with fluorochrome-coupled antibodies specific to CD11b and Gr1 (eBioscience, Carlsbad, CA, USA), whereas the human MDSC were stained with fluorochrome-coupled antibodies specific to CD11b, CD33 and HLADR (eBioscience) for thirty minutes at 4°C. After washing twice with PBS, the cells were analysed using a FACSVerse (BD Biosciences, Billerica, MA, USA).

### MDSC Suppression Assay

MDSC were generated from bone marrow progenitors as described previously. CD8^+^ T cells were isolated from spleen of wild-type C57BL/6 mouse using anti-CD8-coated magnetic beads (Miltenyi), incubated with Cell Trace Violet (CTV; Thermo, Waltham, MA, USA) for 20 min at 37 °C and then washed twice with PBS. Subsequently the CD8^+^ cells were co-cultured with MDSC (3×10^4^ CD8^+^ T cells, 1×10^4^ MDSC cells, 200 μL/well) in 96-well plates coated with 2 μg/mL anti-CD28 and anti-CD3 antibody (eBioscience). After forty-eight hours, the concentration of IFN-γ in the supernatant was detected using the IFN-γ ELISA kit (BD Biosciences) according to the manufacturer’s protocol. The proliferation of CD8^+^ cells labelled with CTV was measured using FACS analysis.

### RNA Isolation and qRT-PCR

Total RNA was isolated using RNA MicroPrep (ZYMO research, Irvine, CA, USA) according to the manufacturer’s protocol. Reverse transcription was performed using the High Capacity cDNA Reverse Transcriptase kit (Applied Biosystems, Foster, CA, USA). Quantitative real-time PCR was performed in a CFX96 Real-Time PCR system (Bio-Rad) using SYBR green qPCR Mix (TaKaRa Bio Inc., Otsu, Shiga, Japan). Expression level of miRNA was normalised to U6 small nuclear RNA, whereas other genes (mRNA) were normalised to β-actin. Primer sequences are specified in Supplementary Table 1.

### MicroRNA Deep Sequencing and Data Analysis

RNA samples from LLC-Exo or CT26-Exo of three independent experiments were pre-mixed with equal qualities. The subsequent sequencing and data analysis was completed at Ribo Bio. in Guangzhou, China. Briefly, Libraries of small RNA from LLC-Exo and CT26-Exo were constructed using TruSeq Small RNA Library Preparation Kits (Illumina, San Diego, CA, USA) according to the manufacturer’s protocol. The libraries were sequenced by Illumina HiSeq2000.

The sequencing read was identical to annotated miRNA sequences in miRBase version 18, only allowing −2 or +2 nt to be template by the corresponding genomic sequence at the 3’ end. The miRNA expression levels were determined on the basis of their read number. Pairwise comparison was performed to identify the significant changes in the miRNA profiles between groups.

### Mimic

A miR-21a-5p mimic and control mimic were purchased from Ribo Bio. in Guangzhou, China.

### Lentivirus Generation and Transfection

Lentivirus generation protocol was mainly based on previous methods(*Nasri et al., 2014*). Briefly, the whole sequence encoding for PDCD4 CDS was synthesised and cloned into lentivirus vector FtetUGW-T. The plasmid was provided by Professor Wan of the Army Medical University in China and by TsingKe Ltd (TsingKe Ltd, Beijing, China). Subsequently, this vector plasmid, along with packaging plasmids psPAX2 and envelope plasmid pMD2.G (4:2:1 ratio)) were co-transfected into the packaging cell line (293T) using lipo 3000 (Thermo) according to the manufacturer’s protocol. After three days, a PDCD4 expressing lentivirus (lv-PDCD4) was harvested. Similarly, the PDCD4 shRNA oligonucleotides (5‘-CTGGACAGGTGTATGATGTGG-3’) were synthesised and cloned into the MISSION® TRC2 pLKO.5-puro Empty Vector (SHC201, Sigma) by TsingKe Ltd. A PDCD4 shRNA expression lentivirus (lv-shPDCD4) was generated as described previously. The viral suspensions lv-PDCD4 or lv-shPDCD4 were pre-mixed with Polybrene (Sigma-Aldrich, St. Louis, MO, USA) to a final concentration of 2 mg/mL and then used for infection of the BM cells. Infection was performed in 24-well plates (5×10^5^ cells/mL) with an eight-hour incubation at 37°C and 5% CO2. Then cells were washed and re-suspended in the MDSC induction medium with or without TEXs.

### CRISPR/Cas9

For knocking-out miR-21a in LLC cells, a target guide RNA sequence was designed as miR21a-gRNA: TGATAAGCTATCCGACAAGG. Double strands of miR21a-gRNA were synthesised and cloned to LentiCRISPR-V2 plasmid (Addgene, Cambridge, MA, USA) by TsingKe Ltd. One-week after transfection of the LentiCRISPR-V2 virus, monoclonal LLC cell was sorted into 96-well plates by FACS. To identify the miR-21a-knockout LLC cell clones, genomic DNA PCR was performed (using primers of miR21a-gR-F/miR21a-gR-R, as indicated in Supplementary Table 2) and the PCR products were subjected to DNA sequencing verification. Subsequently miR-21a expression was determined using qTR-PCR as described previously.

### Luciferase Reporter Assay

Full lengths of the PDCD4-WT-UTR and PDCD4-MT-UTR (Supplementary Table 3) were synthesised and cloned into psiCHECK-2 vector by TsingKe Ltd. respectively. The 293T cells were co-transfected with 500 ng psiCHECK-2–PDCD4-3ʹUTR (wild type or mutant type) and 50 nM miR-21 mimic or control mimic using TransFECT (Ribo Bio) according to the manufacturer’s instructions. After forty-eight hours, the Luciferase activities of whole cell lysate were measured using Dual-Luciferase Reporter Assay System (Promega, Madison, WI, USA) by a GloMax20/20 Luminometer (Promega). Luciferase activity was normalised by Renilla/Firefly luciferase signal in 293T cells.

### Western Blotting

Proteins were extracted from cells and exosomes using RIPA Lysis (Thermo) containing Protease/Phosphatase Inhibitor Cocktail (Cell Signalling Tech, Denver, MA, USA). Samples were normalised according to BCA protein assay kit (Thermo Scientific Pierce) and proteins were separated following an electrophoretic gradient across polyacrylamide gel electrophoresis (PAGE) and then blotted onto polyvinylidene fluoride (PVDF) membranes (Millipore Corp, billerica, MA, USA). The PVDF membranes were blocked with 5% w/v Bovine Serum Albumin (BSA; Sigma)for one hour at room temperature and incubated overnight at 4 °C with the following primary antibodies: anti-CD63 (BioLegend, San Diego, CA, USA), anti-Cytochrome c, anti-HSP90, anti-STAT3, anti-pSTAT3 (Cell Signaling), anti-HSP72 (Enzo Life Sciences, Farmingdale, NY, USA), anti-PDCD4, anti-β-actin (Proteintech). The membranes were then incubated with secondary antibodies (HRP-linked anti-mouse IgG; Sigma) for forty minutes at room temperature. PVDF membranes were washed with TBS-Tween20 three times at ten minute intervals and digital images were obtained using the ChemiDoc^™^ Touch Imaging System (BioRad) with a Pierce^™^ ECL Western Blotting Substrate (Thermo).

### Enzyme-linked Immunosorbent Assay

ELISA kits were used to detect IL-6, IL-10, IFN-γ (BD Biosciences), VEGF and TNF-α (R&D Systems, Minneapolis, MN, USA) according to the manufacturer’s protocol.

### Statistics

Results were analysed with GraphPad Prism 5. Any means of continuous outcome variables were tested with a two-tailed student t test. P < 0.05 was considered statistically significant. Results are expressed as means ± SEM.

## Acknowledgements

We thank Professor Ying Wan for providing LLC-luc (an orthotopic luciferase stably expressed LLC) cells, lentivirus vector FtetUGW-T and LentiCRISPR-V2 plasmid from the Army Medical University of China.

## Additional information

### Funding

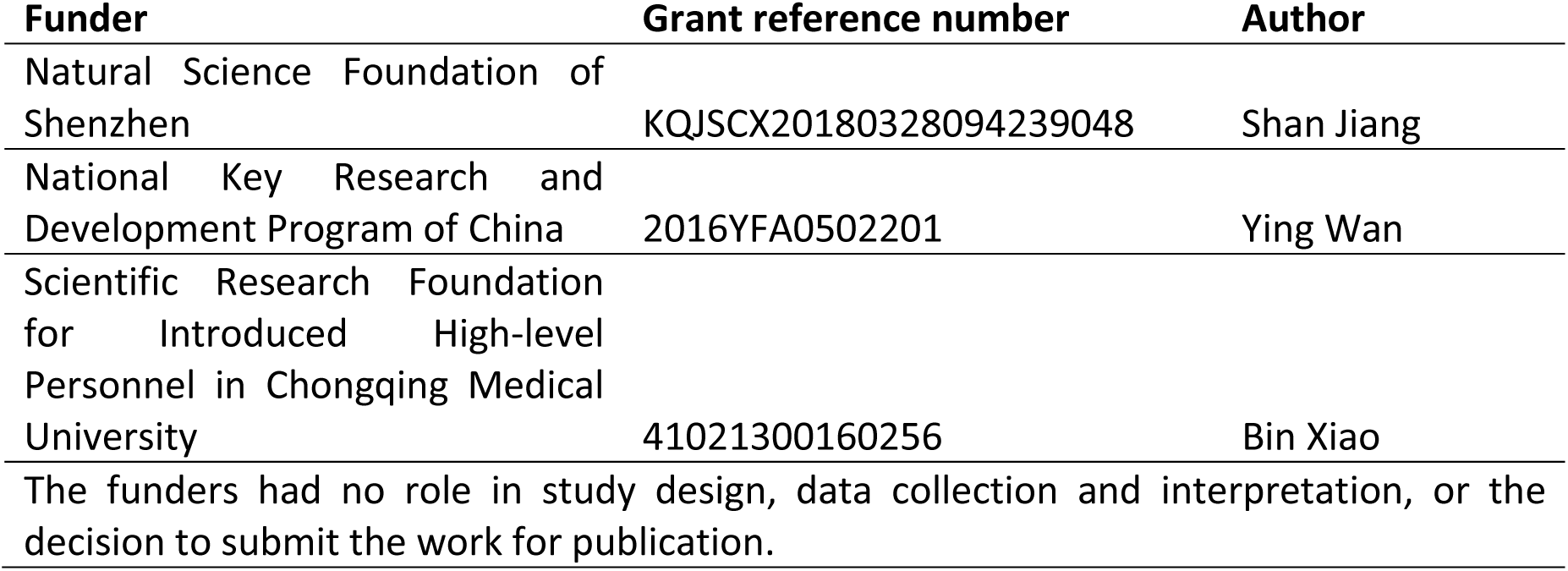

### Author contributions

S.J. and X.Z. conceived and designed the work. X.Z., F.L and J.Z. collected and analyzed the data for this work. Y.W., B.X., X.Z., F.L., Y.T. and J.Z. participated in the preparation of the manuscript. J.S. and X.Z. wrote the manuscript that was reviewed by all authors.

### Author ORCIDs

Shan Jiang: https://orcid.org/0000-0002-8297-0099

### Ethics

Animal experimentation: Animal studies were carried out according to protocols approved by the Medicine Animal Care Committee of the Army Medical University (Chongqing, China). Number of animals for study and unnecessary suffering was minimized as much as possible.

## Supplementary Figures

**Supplementary Figure 1.**
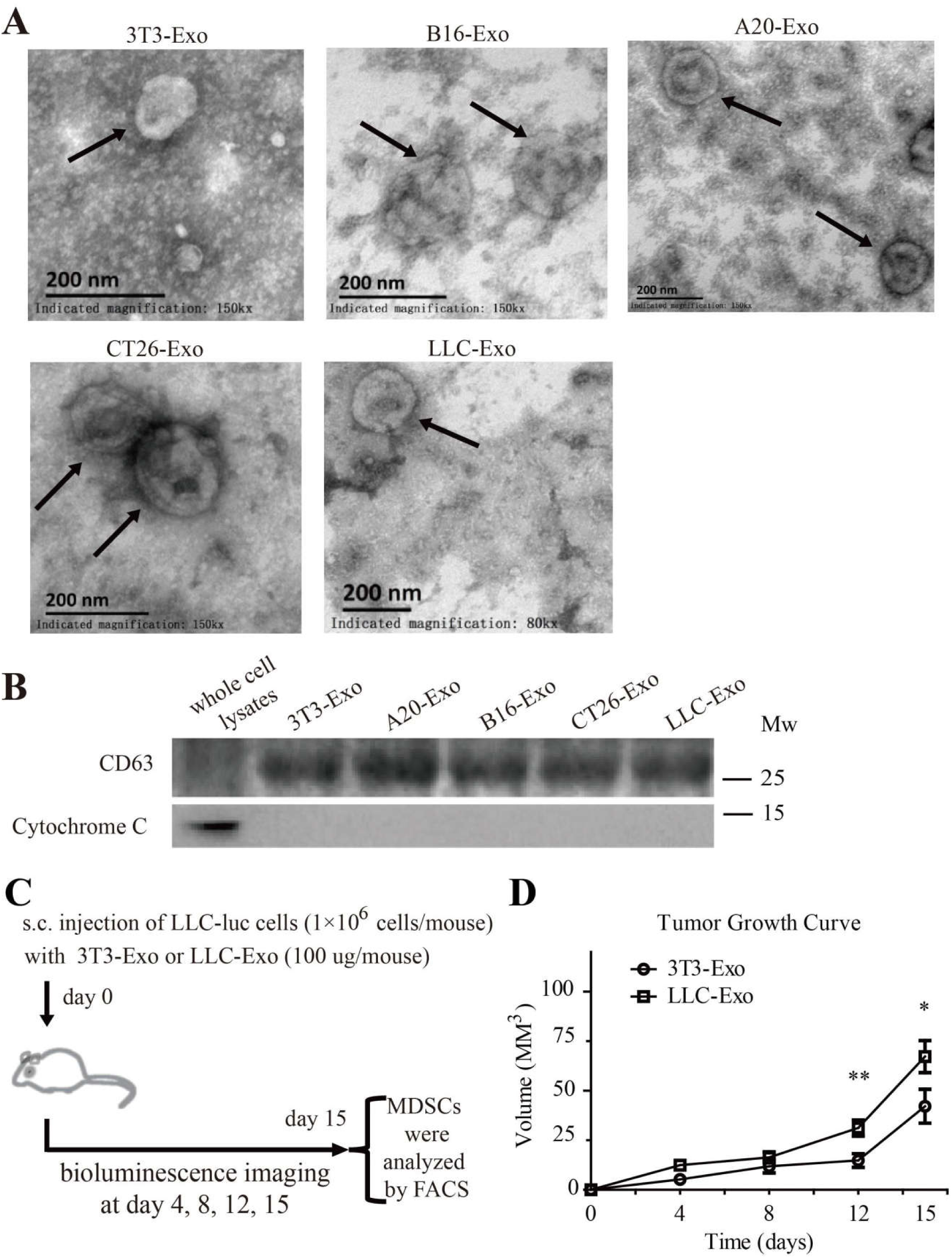
Exosomes isolated from 3T3, A20, B16, CT26 and LLC cells. (**A**) Exosomes were stained with uranyl acetate and then visualised using a Philips-FEI TecnaiT10 TEM. Scale bar = 200 nm. (**B**) CD63 and Cytochrome C expression in whole cell lysates of 3T3 cell and exosomes isolated from different cells. (**C**) Schematic description of the *in-vivo* experimental design. (**D**) LLC cells with luciferase expression (1×10^6^ cells in 100 μL PBS per mouse) mixed with exosomes (100 μg in 100 μL PBS per mouse) produced by 3T3 cells (3T3-Exo) or LLC cells (LLC-Exo) were subcutaneously injected into the C57BL/6 mouse. Each group contained eight animals. The tumour length and width were measured every three days over fifteen days. Tumour volume was calculated with formula (V = Length × Width^2^/2). A two-tailed student t test was used and error bars represent the mean ± sem.

**Supplementary Figure 2.**
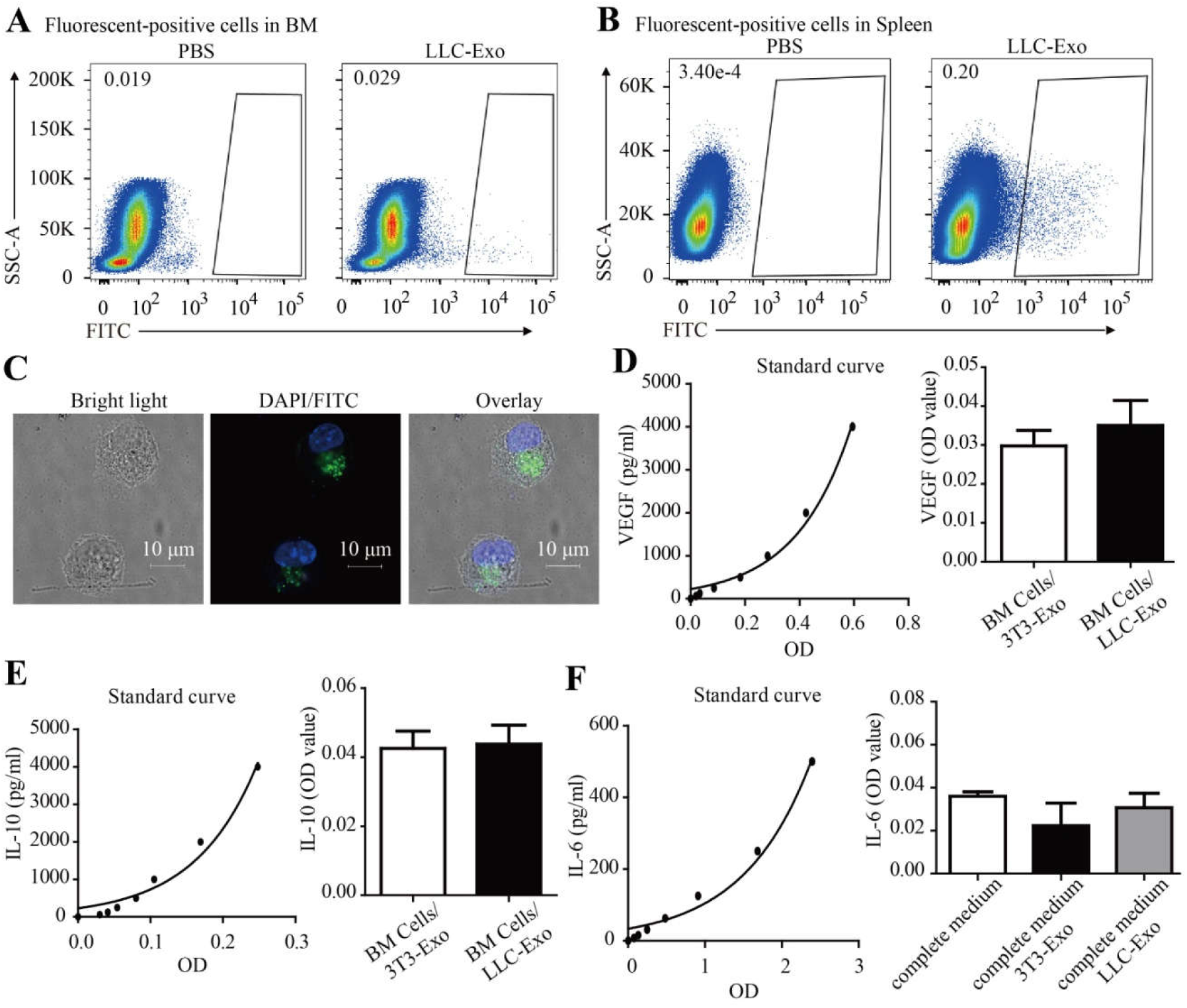
(**A-B**) C57BL/6 mice were injected intravenously with PKH67-labeled LLC-Exo (100 μg in 100 μL PBS per mouse) or PBS. Twenty-four hours later, mice were sacrificed. The BM (**A**) and spleen (**B**) cells were recovered and subjected to FACS analysis for PHK67 (FITC) positive cells. (**C**) LLC-Exo (50 μg/mL) were labeled with green lipophilic fluorescent dye PKH67 and incubated with BM cells (5×10^5^ cells/ mL). Four to Five hours later, the uptake of exosomes was detected by confocal laser scanning microscopy, scale bar = 10 μm. (**D-E**) Mice BM cells were isolated and cultured in an MDSC-inducing medium with LLC-Exo or 3T3-Exo (50 μg/mL). After culture for seven days, the supernatants of cell cultures were collected and the concentration of VEGF, IL-10 was determined by ELISA. (**F**) The LLC-Exo and control 3T3-Exo were added into an MDSC-inducing medium (50 μg/mL) for two days, the IL-6 concentration in the supernatantswas determined by ELISA. Error bars represent the mean ± sem.

**Supplementary Figure 3.**
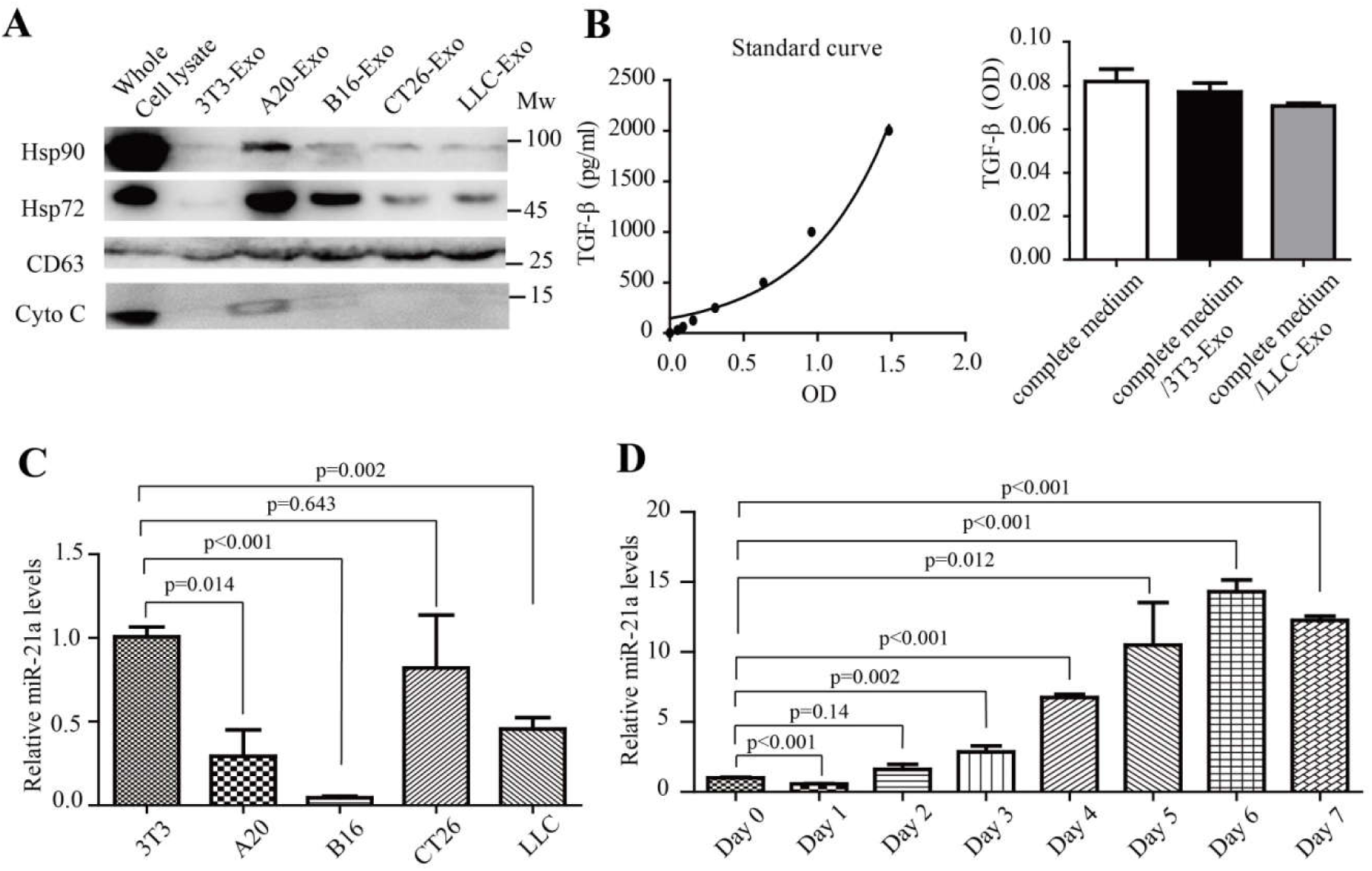
(**A**) The expression of heat-shock protein HSP90 and HSP72 in exosomes of 3T3-Exo, A20, B16-Exo, CT26-Exo and LLC-Exo and whole cell lysate of 3T3 cells were examined using Western blotting. (**B**) The concentration of TGF-β in LLC-Exo and 3T3-Exo was determined using ELISA. (**C**) Total RNAs were extracted from 3T3, A20, B16, CT26 and LLC cells. The amount of miR-21a in these cells was determined by quantitative RT-PCR. (**D**) Mice BM cells were isolated and cultured (5×10^5^ cells/mL) in an MDSC-inducing medium. After seven days, total RNAs were extracted from these BM cells each day. Then the amount of miR-21a was determined using quantitative RT-PCR. A two-tailed student t test was used and error bars represent the mean ± sem.

**Supplementary Figure 4.**
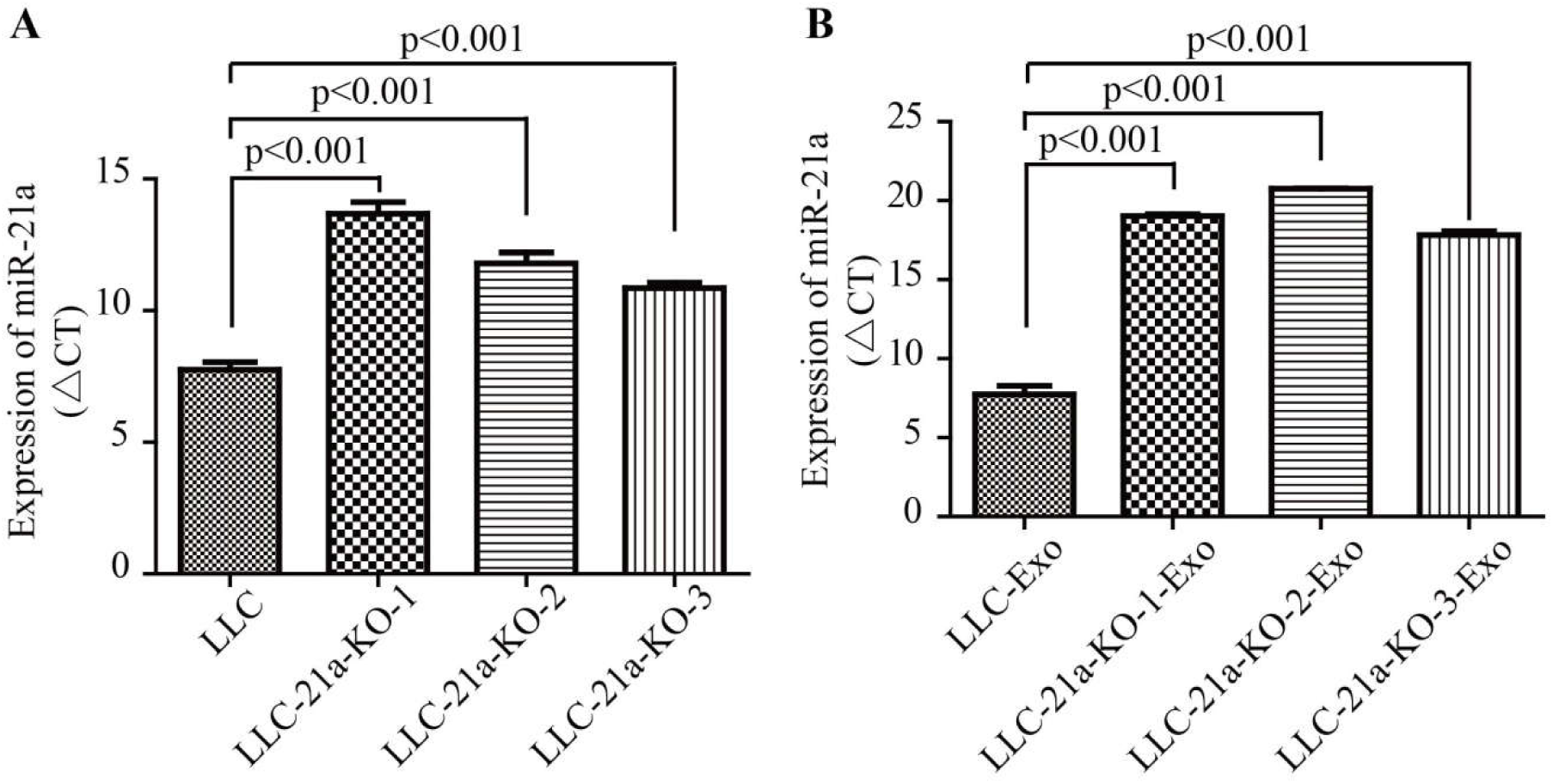
Total RNA was isolated from single clones of LLC-21a-KO cells and the parental LLC cells. (**A**) The amount of miR-21a in these cells was determined by quantitative RT-PCR (**B**) The amount of miR-21a in exosomes was determined by quantitative RT-PCR. LLC-21a-KO-1, LLC-21a-KO-2, LLC-21a-KO-3 were three single clones from LLC-21a-KO cells. A two-tailed student t test was used and error bars represent the mean ± sem.

**Supplementary Figure 5.**
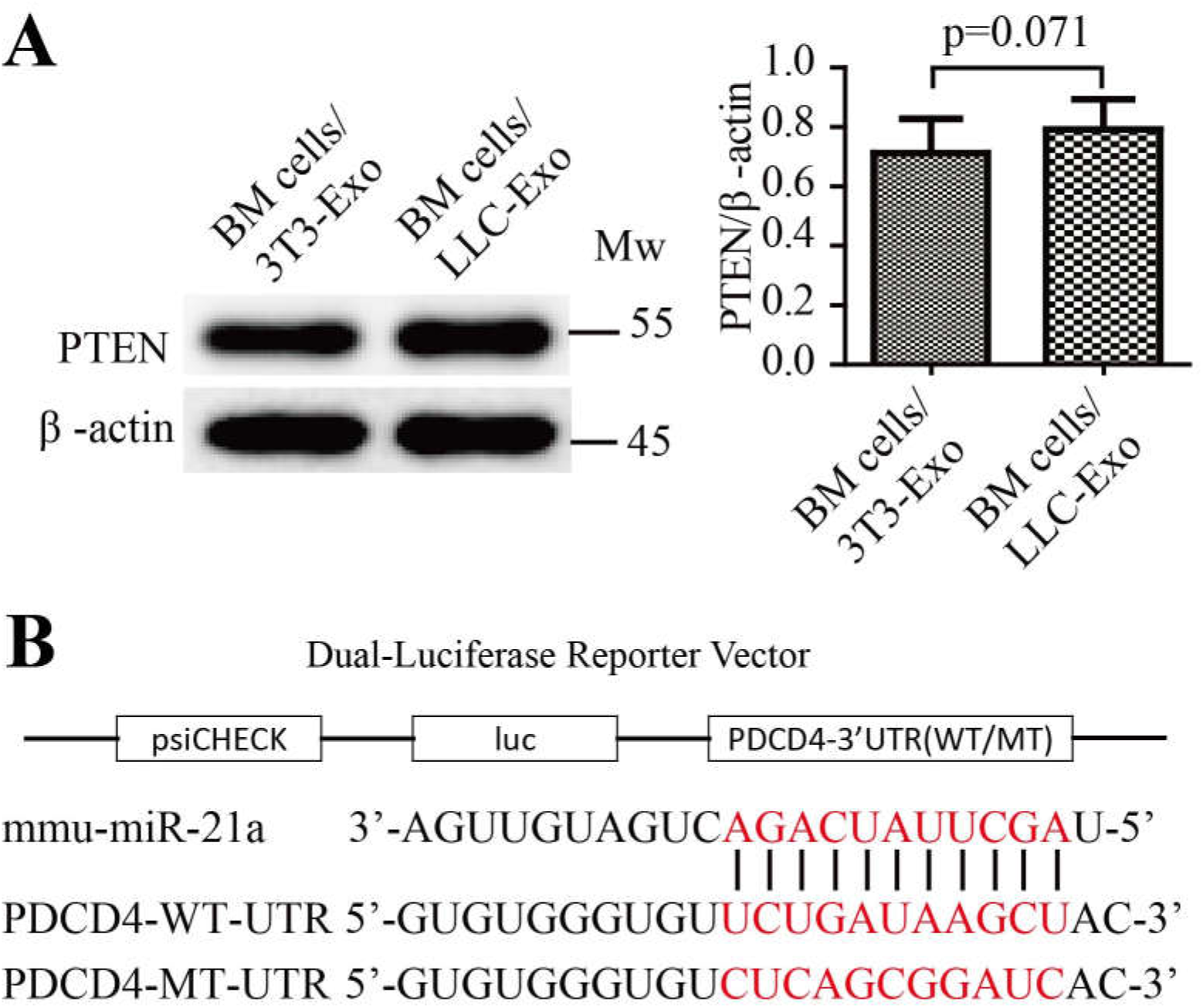
(**A**) Mice BM cells were isolated and cultured (5×10^5^ cells/mL) in an MDSC-inducing medium with 3T3-Exo or LLC-Exo (50 μg/mL) for seven days. The amount of PTEN in BM cells was determined by western blotting. Experiments were repeated for three times and a two-tailed student t test was used. Error-bars represent the mean ± sem. (**B**) Normal and mutated mRNA sequence in the 3′-UTR of PDCD4 predicted to be bound by miR-21a. The seed-binding site in the 3′-UTR of PDCD4 mRNA was highlighted in red colour.

**Supplementary Figure 6.**
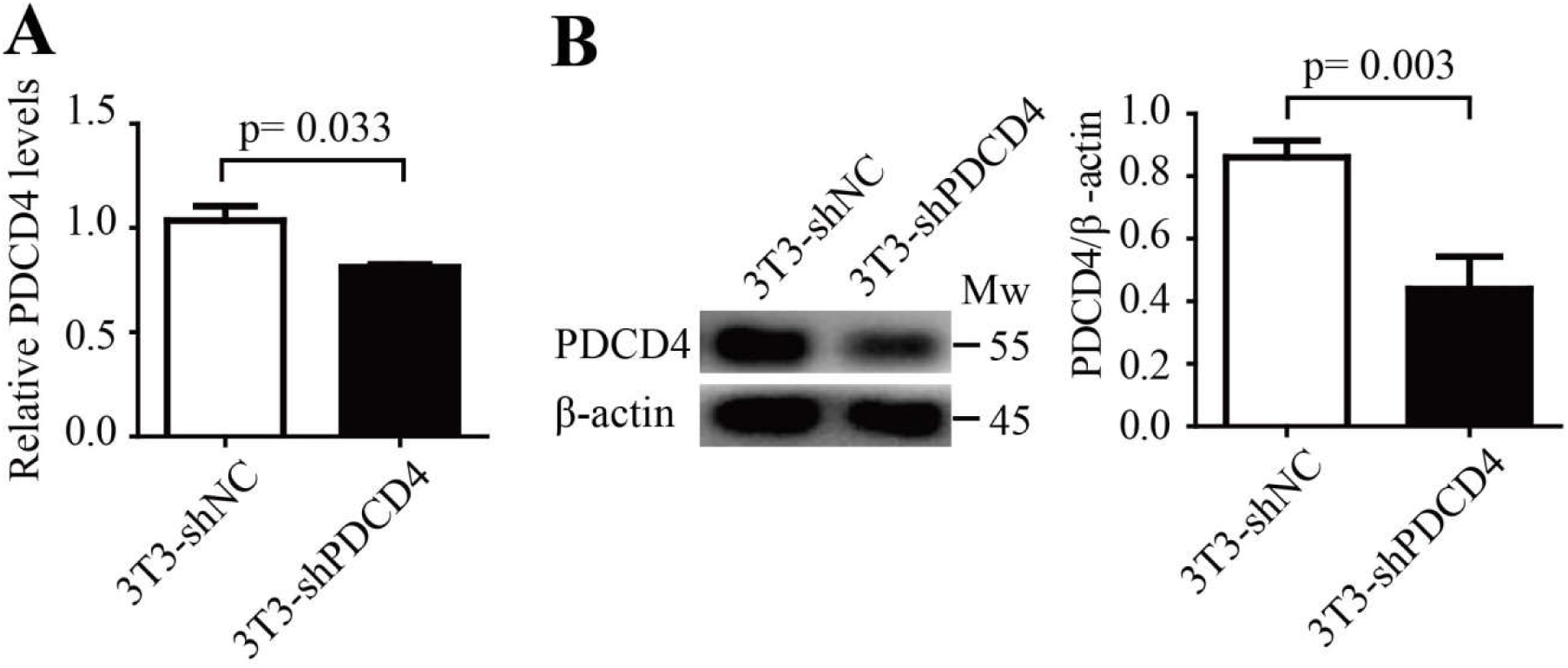
The 3T3 cells were infected with lentivirus (lv-shNC or lv-shPDCD4) GFP-positive 3T3 cells were sorted out using FACS. Then total RNA and whole cell lysates were harvested. (**A**) The amount of PDCD4 mRNA was determined using quantitative RT-PCR. (**B**) The amount of PDCD4 protein was determined using Western blotting. A two-tailed student t test was used and error bars represent the mean ± sem.

**Supplementary Figure 7.**
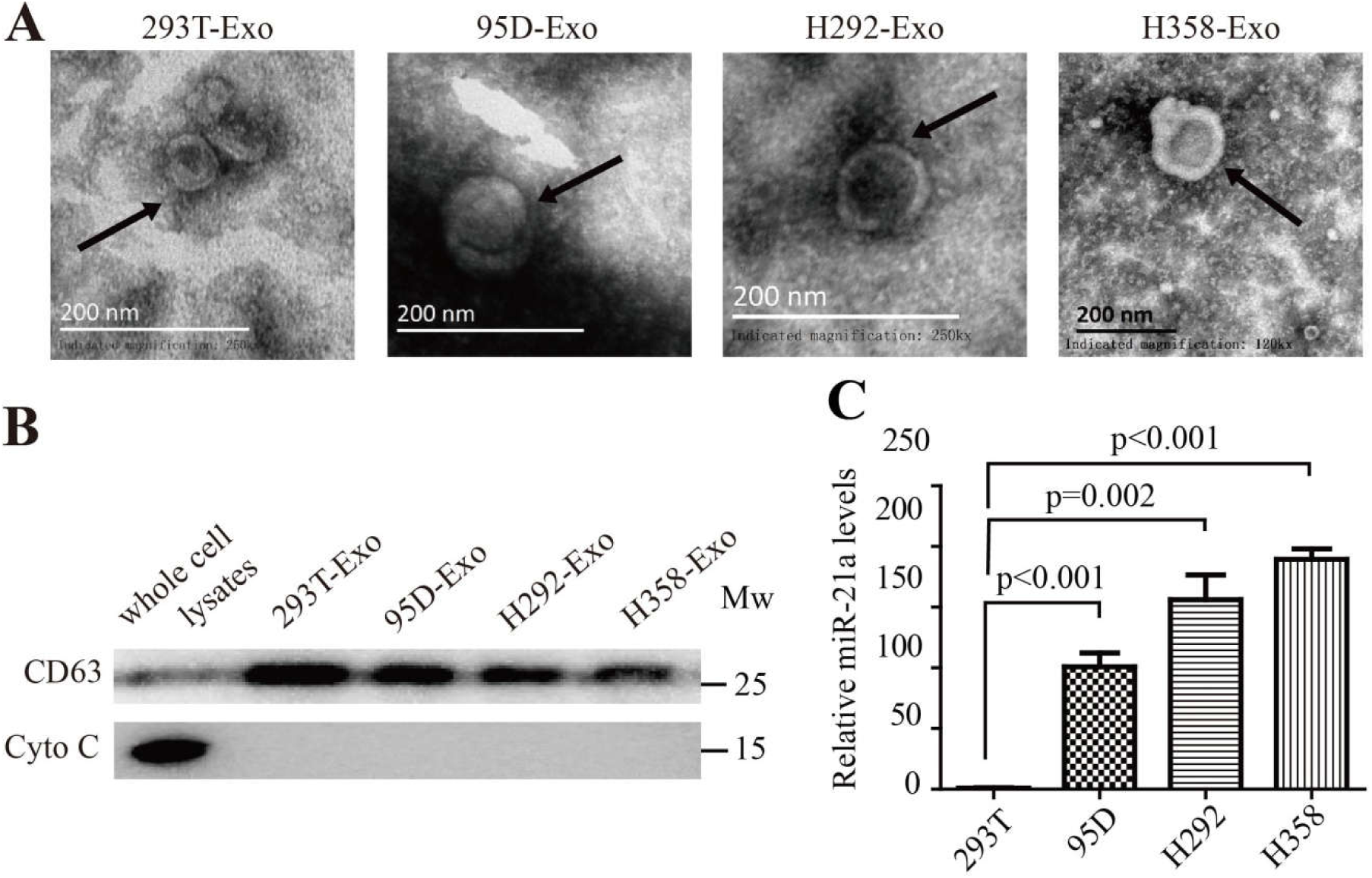
(**A**) Exosomes were isolated from human lung cancer cells and then observed by Electron microscope. Scale bar, 200 nm. (**B**) The expression of CD63 and Cytochrome C in the whole cell lysates or exosomes of 293T cells, 95D cells, H292 cells and H358 cells were determined by Western blotting. (**C**) The amount of miR-21a in 293T cells, 95D cells, H292 cells and H358 cells were determined by quantitative RT-PCR. A two-tailed student t test was used and error bars represent the mean ± sem.

**Supplementary Table 1.**
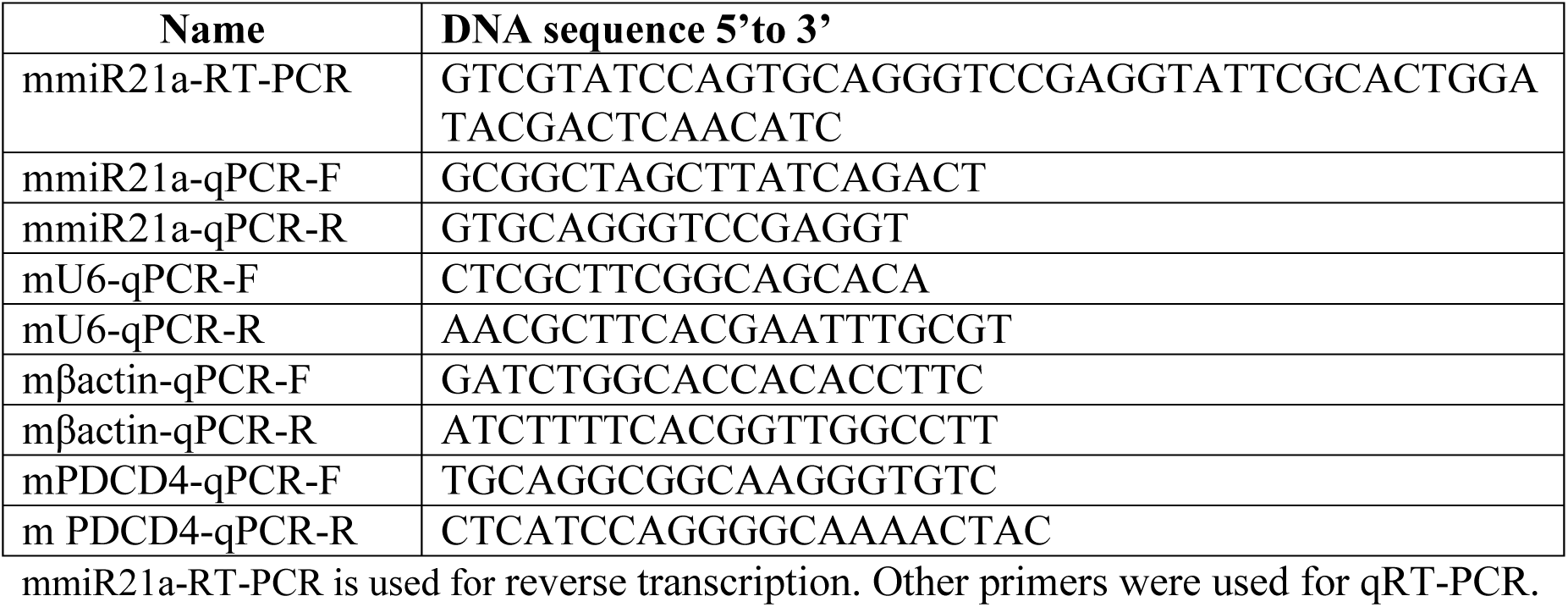
A list of primers used in this study.

**Supplementary Table 2.**
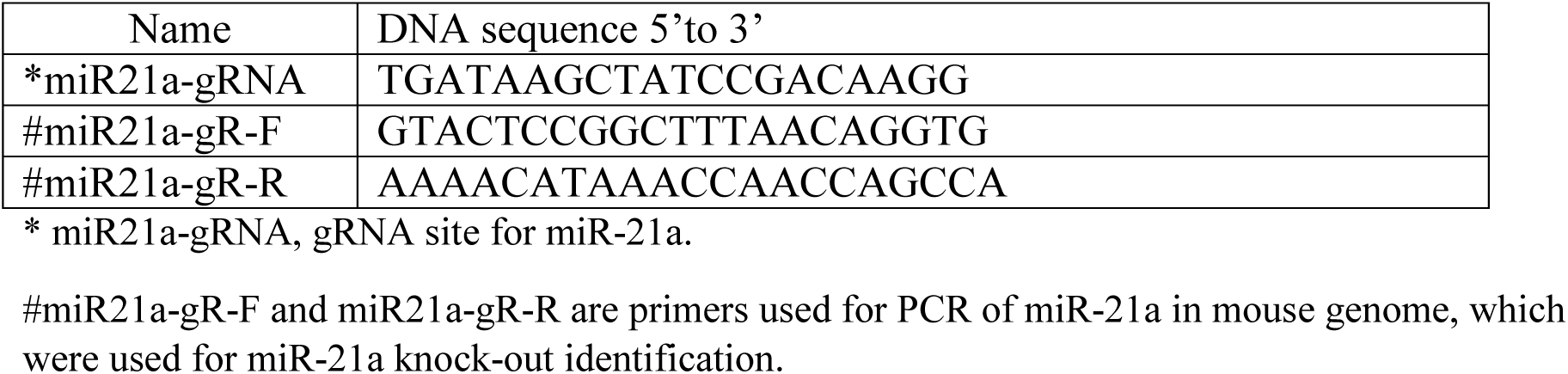
The DNA sequence of guide-RNA used to CRISPR-Cas9 system in this study.

**Supplementary Table 3.**
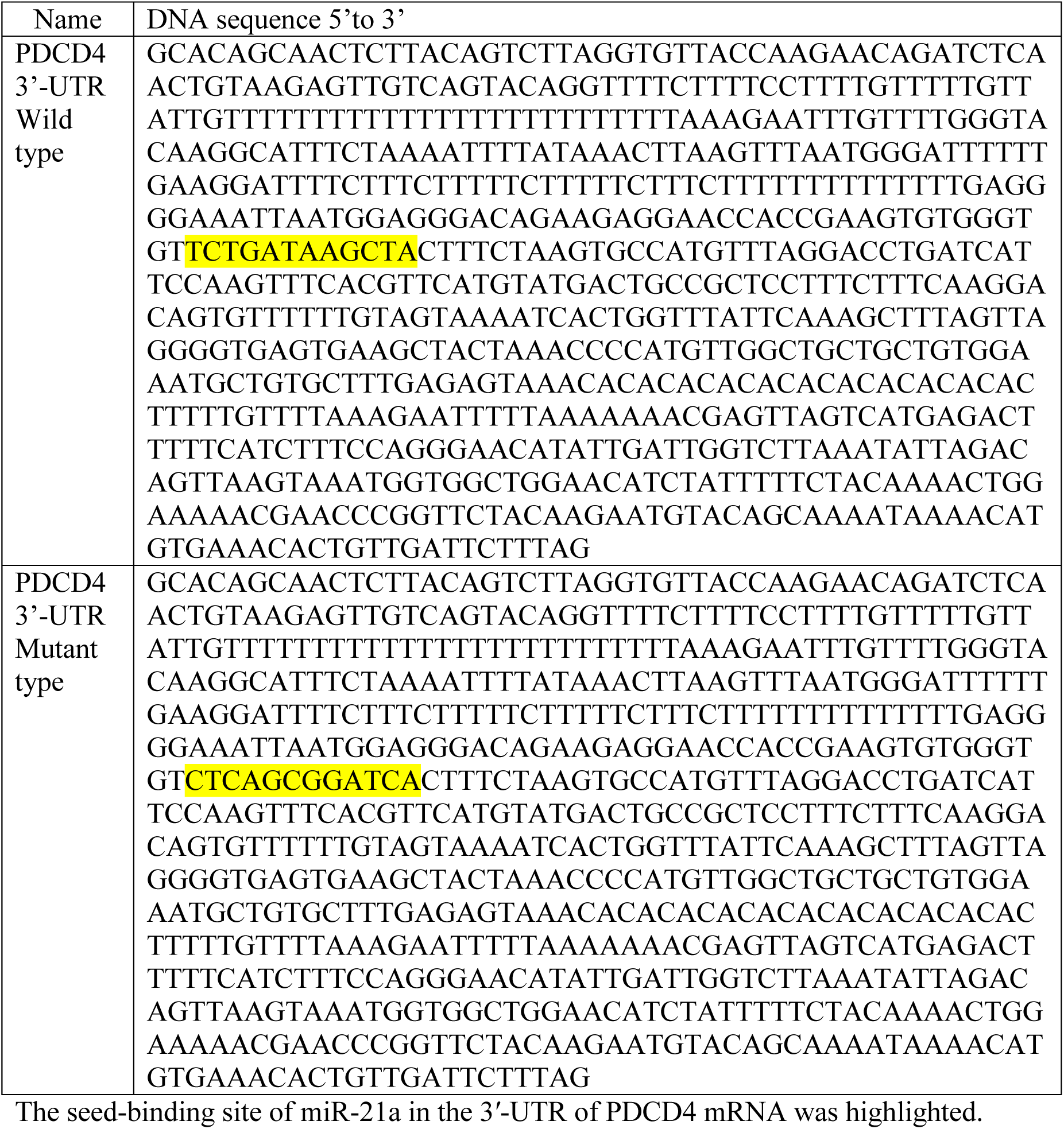
The DNA sequences of the wild type and mutant type of 3’-UTR of PDCD4 gene that was cloned into the psiCHECK-2 vector.

